# Assessing the importance of resistance, persistence and hyper-mutation for antibiotic treatment success with stochastic modelling

**DOI:** 10.1101/2022.04.07.487440

**Authors:** Christopher Witzany, Roland R. Regoes, Claudia Igler

## Abstract

Antimicrobial resistance poses a rising threat to global health, making it crucial to understand the routes of bacterial survival during antimicrobial treatments. Treatment failure can result from genetic or phenotypic mechanisms, which diminish the effect of antibiotics. By assembling empirical data, we find that, for example, *Pseudomonas aeruginosa* infections in cystic fibrosis patients frequently contain persisters, transiently non-growing and antibiotic-refractory subpopulations, and hyper-mutators, mutants with elevated mutation rates and thus higher probability of genetic resistance emergence. Resistance, persistence and hyper-mutation dynamics are difficult to disentangle experimentally. Hence, we use stochastic population modelling and deterministic fitness calculations of bacterial evolution under antibiotic treatment to investigate how genetic resistance and phenotypic mechanisms affect treatment success. We find that treatment failure is caused by resistant mutants at lower antibiotic concentrations (with high final bacterial numbers), but by persistence phenotypes at higher antibiotic concentrations (with low final bacterial numbers). Facilitation of resistance occurs through hyper-mutators during treatment, but through persistence only after treatment is discontinued, which allows for persisters to resume growth and evolve resistance in the absence of antibiotics. Our findings highlight the time- and concentration-dependence of different bacterial mechanisms to escape antibiotic killing, which should be considered when designing ‘resistance-proof’ antimicrobial treatments.

## Introduction

The evolution of antimicrobial resistance is a global and growing threat to human lives and contemporary medicine (Murray et al., 2022). Studies have shown that antibiotic (AB) resistance is a complicated trait that can be facilitated by resistance-enabling mechanisms, such as persistence and hyper-mutation (Levin-Reisman et al., 2017; Levin-Reisman et al., 2019; Mehta et al., 2019; Rodriguez-Rojas et al., 2021). Therefore, to ensure prolonged efficacy of current and future ABs, it is crucial to investigate how resistance-enabling mechanisms impact the emergence of resistance and treatment failure in general.

In long-lasting infections, such as those caused by *Pseudomonas aeruginosa* or *Mycobacterium tuberculosis*, genetic resistance can emerge via random chromosomal mutations over the course of treatment and cause complications or treatment failure (Oliver et al., 2000; Castro et al., 2021). The speed by which mutations arise is hence crucial for pathogen survival. This mutation rate is heavily influenced by replication errors and can be increased about 100 to 1000-fold (Mena et al., 2008; Lee et al., 2012) in mutants that have faulty replication pathways, so-called hyper-mutators. Most mutations will be deleterious and decrease the fitness of hyper-mutators. However, hyper-mutators are known to flourish in highly fluctuating environments by acquiring beneficial mutations, like AB resistance, which can outweigh the cost of deleterious mutations (Giraud et al., 2002; Travis & Travis, 2002; Mena et al., 2008). Hyper-mutators thereby pose a considerable threat to the efficacy of ABs by significantly increasing the probability of resistance emergence (Figure 1A), especially, since empirical studies suggest their prevalence in chronic infections with *P. aeruginosa* (Figure 1B, Text S1), *Escherichia coli* (Labat et al., 2005) and *Staphylococcus aureus* (Prunier et al., 2003).

**Figure 1.**
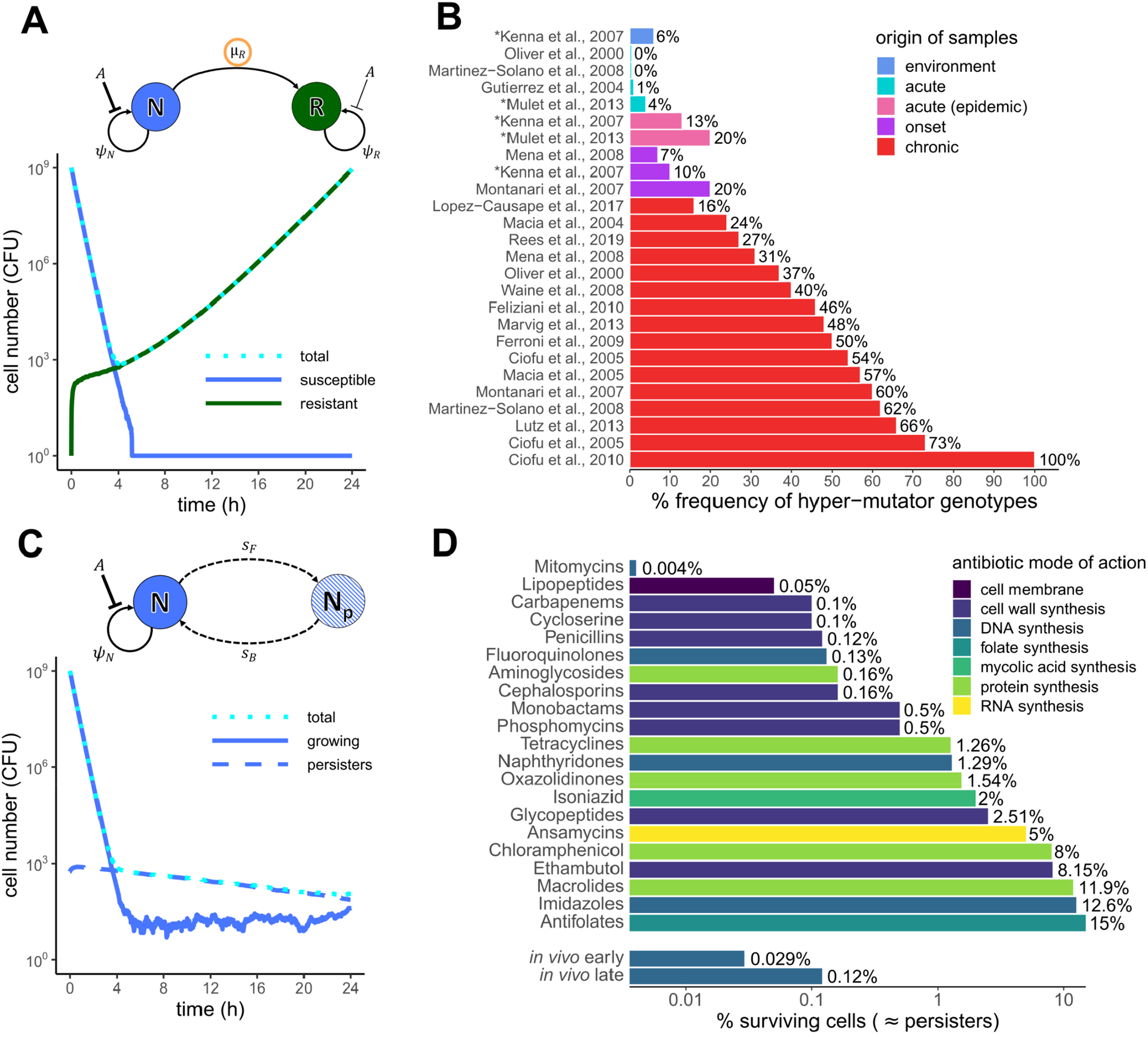
Antibiotic treatment failure via genetic resistance and phenotypic persistence. **A**) Population dynamics of *de novo* resistance evolution over the course of a one-time AB treatment obtained by stochastic modelling. Shown are the number of total bacterial cells (CFU) as the cyan, dotted line, the susceptible starting population (N) as the solid blue line and the emerging resistant population (R) as the solid green line. Growth and growth inhibition by ABs (*A*) are modelled as separate stochastic processes (Methods), but for simplicity shown here as net growth rate *ψ*_*N*_ for susceptibles (as given by MIC). Resistants (R) emerge from N by mutation at rate µ_R_, which would be increased for hyper-mutators. R grows at *ψ*_*R*_, which is given by the cost of resistance and its lower vulnerability to ABs (MIC_R_=10xMIC). AB treatment for **A**) and **C**) was simulated with an AB dose of 10xMIC for 24h (Methods, Table S1). **B**) Hyper-mutator frequencies of *Pseudomonas aeruginosa* (%) compiled from empirical studies. Shown are frequencies for samples from the environment, or from patients with regular acute infection, epidemic infection, onset of chronic infection and chronic infection. For studies marked with an asterisk (*) isolate level data is shown as patient level data was not available (Text S1). **C**) Characteristic biphasic killing curve obtained from a two-state population model with switching (*s*_*F*_, *s*_*B*_) between growing (N) and persister state (N_p_) based on the model by Balaban et al. (2004) (adapted to our notation). The persister subpopulation (N_p_) is shown as the blue, dashed line, other colours as in **A. D**) Persister numbers of multiple species as % of cells surviving exposure to different AB classes (colour-coded according to mode of action) from *in vitro* and *in vivo* studies assembled from literature (Salcedo-Sora and Kell (2020), Mulcahy et al. (2010); Text S1). For comparison with clinical data, *in vivo* persister numbers from isolates from early and late stages of a chronic infection with *P. aeruginosa* are shown separately.

While emergence of genetic resistance is still considered the main cause of treatment failure, it is becoming increasingly clear that non-genetic, transient mechanisms also enable bacteria to survive AB treatment. One such mechanism is persistence, which describes a phenotypic state, defined by the formation of bacterial subpopulations that are in a temporary non-growing state, which allows them to be transiently refractory, i.e. unaffected by ABs (Balaban et al., 2019). This can be observed as a biphasic killing curve in the presence of ABs (Figure 1C), where the growing population dies rapidly, leaving the smaller persister population, which declines at a much slower rate. Persistence has been reported for many antibiotic classes (Figure 1D, Text S1) and has been found to facilitate the evolution of resistance. This facilitation occurs due to higher and prolonged survival of susceptible bacteria, thereby increasing the opportunity for mutations to occur (Levin-Reisman et al., 2017) – as opposed to increasing the mutation rate itself. Moreover, there are known mutants that generate larger persister subpopulations than the wildtype, so-called high-persisters (Moyed & Bertrand, 1983; Wolfson et al., 1990; Balaban et al., 2004). High-persistence mutations, and persistence in general, are beneficial in highly fluctuating environments (Kussell et al., 2005; Van den Bergh et al., 2016) and are frequent in chronic infections with *E. coli* (Schumacher et al., 2015), *Candida albicans* (Lafleur et al., 2010) and *P. aeruginosa* (Bartell et al., 2020). Notably, for cystic fibrosis (CF) patients, persister subpopulations increase over the course of chronic infection (Figure 1D). This is likely due to the emergence of high-persistence mutants, which, like hyper-mutators, have been reported more frequently at later time points of infection (Mulcahy et al., 2010) (Figure 1B). Therefore, high-persisters could provide a pool of genetically susceptible, but viable cells, that survive AB treatment and thereby cause the “paradox of chronic infections”, which describes the phenomenon of chronically recalcitrant infections with non-resistant pathogens (Lewis, 2010).

The involvement of hyper-mutation and high-persistence mutations in treatment failure of chronic infections suggests that genetic and phenotypic mechanisms are not only both beneficial for survival in changing environments (such as AB treatment), but also that they might interact with one another. Indeed, hyper-mutation and high-persistence mutations have been found in the same clinical strain of *P. aeruginosa* (Mulcahy et al., 2010), a combination that we will refer to as a mutator-persister. The presence of mutator-persisters in chronic infections suggests the possibility that hyper-mutators could lead to a beneficial high-persistence mutation. However, it could also be the other way around, with the fitter high-persisters causing sufficient population survival to enable the evolution of less fit hyper-mutators. Interestingly, Mulcahy et al. (2010) found mutator-persisters to be also genetically resistant against ABs, indicating that resistance might still be beneficial, and potentially facilitated by the presence of both, hyper-mutation and high-persistence.

Disentangling the contributions of genetic and phenotypic mechanisms to treatment failure poses several challenges. Examining the dynamics of persistence, and especially high-persistence, is inherently difficult experimentally due to the stochastic and phenotypic nature of this trait (Kim & Wood, 2016; Balaban et al., 2019). Including hyper-mutators will likely aggravate this problem, for example due to the necessity of more experimental replicates owing to increased stochasticity (Raynes & Weinreich, 2019) and the higher number of different mutations that can arise. Here we use stochastic modelling to investigate (a) how mutant populations of hyper-mutators (M), high-persisters (P) and resistant (R) cells – as well as all their respective combinations – evolve over time under different AB concentrations, (b) how they affect treatment outcome and (c) derive analytical calculations to understand the simulation outputs through the long-term fitness of specific genotypes under AB treatment. We show that R, M and P populations cause or facilitate treatment failure at distinct AB concentrations, infection time scales and final cell numbers. Our goal is not to make precise quantitative predictions, but rather utilize mathematical modelling to explore under which conditions these populations can or should evolve.

## Results

### Modelling of persistence, mutator and resistance dynamics

To investigate the relative importance of phenotypic and genetic mechanisms of bacterial cells to escape antibiotic killing, we use a stochastic pharmacodynamic model (Figure 2) to simulate population dynamics during antibiotic (AB) treatment of acute infections (Methods, Text S2). Bacterial persistence can complicate treatment by allowing susceptible bacteria to survive inhibitory AB concentrations, without having to acquire genetic changes (Lewis, 2010). To capture this, we first focus on a submodel that describes the susceptible genotype (N), its persister subpopulation (N_p_) and a resistant mutant (N_R_), which cannot switch into a persister state (Figure 2, 3A). N and N_p_ stochastically switch back and forth at rates *s*_*B*_ and *s*_*F*_ (Figure 1B). We start with N and N_p_ in equilibrium according to the stochastic switching in the absence of AB treatment (Methods), i.e. the growing, AB-sensitive subpopulation N at ∼10^9^ CFU and the non-growing, AB-refractory persisters N_p_ at ∼5×10^2^ CFU. From the growing subpopulation (N) a *de novo* resistant genotype (N_R_) with MIC_R_ = 10xMIC of susceptible cells can arise via random mutation (Figure 1D). Since we assume that mutations are linked to cellular growth the non-growing persister subpopulation cannot mutate. Hence, N_p_ can only facilitate resistance emergence through prolonging the survival of the N population (Figure S1). We simulate 8 days of treatment with an AB dosage equal to MIC_R_ given once every 24h, which decays over time (Table S1). As expected, N declines rapidly under treatment and N_p_ declines at a much slower rate, which is determined by the rate of switching back to N, displaying the characteristic biphasic killing curve of persisters (Balaban et al., 2004, Figure 1A, 3A). N_R_ evolves rapidly from the N population and reaches carrying capacity after approximately 2 days. Note that the time until N_R_ reaches carrying capacity is dependent on the cost of resistance. In contrast N and N_p_ are fully eradicated after about 4 days. This demonstrates that while persistence prolongs clearance of the susceptible genotype, the emergence of resistance seems to be the sole cause of treatment failure under the simulated regimen.

**Figure 2.**
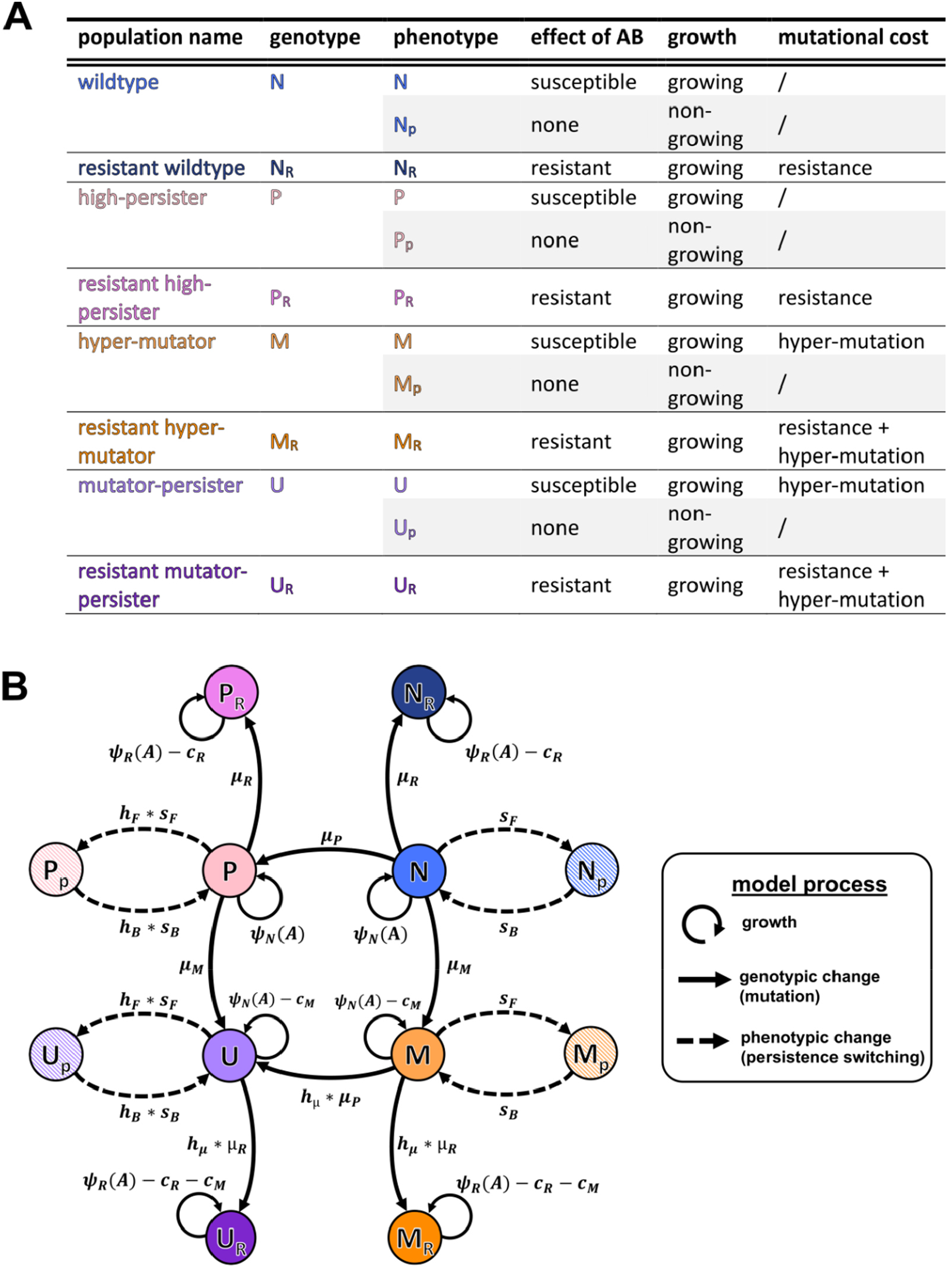
Schematic of the stochastic model describing resistance, high-persistence and hyper-mutator dynamics. **A**) Overview of all modelled genotypes, their phenotypic states (grey background for non-growing phenotypes), susceptibility to antibiotics (ABs), growth state, and incurred cost. **B**) Illustration of the full mathematical model. The eight genotypes consist of the WT (N), hyper-mutators (M), high-persisters (P), mutator-persisters (U) and their corresponding resistant mutants denoted by subscript R. Persister phenotype states (hatched) are denoted by a subscript p. Switching between these two states (dashed arrows) happens at rates *s*_*F*_ and *s*_*B*_ for N and M, and with *h*_*F*_- and *h*_*B*_-fold increase for P and U. Growth rates as determined by AB sensitivity (Eq. 2, Text S1) are given as *ψ*_N_ for susceptible (MIC = 1) and as *ψ*_*R*_ (MIC_R_ = 10xMIC) for resistant genotypes, together with growth costs due to resistance, *C*_*R*_, and due to hyper-mutation, *C*_*M*_. Solid arrows show mutational transitions between genotypes, which happen at rates µ_*M*_, µ_*P*_ and µ_*R*_. Mutators (M, U) have *h*_µ_-fold increased mutation rates. See Table S1 for parameter values.

**Figure 3.**
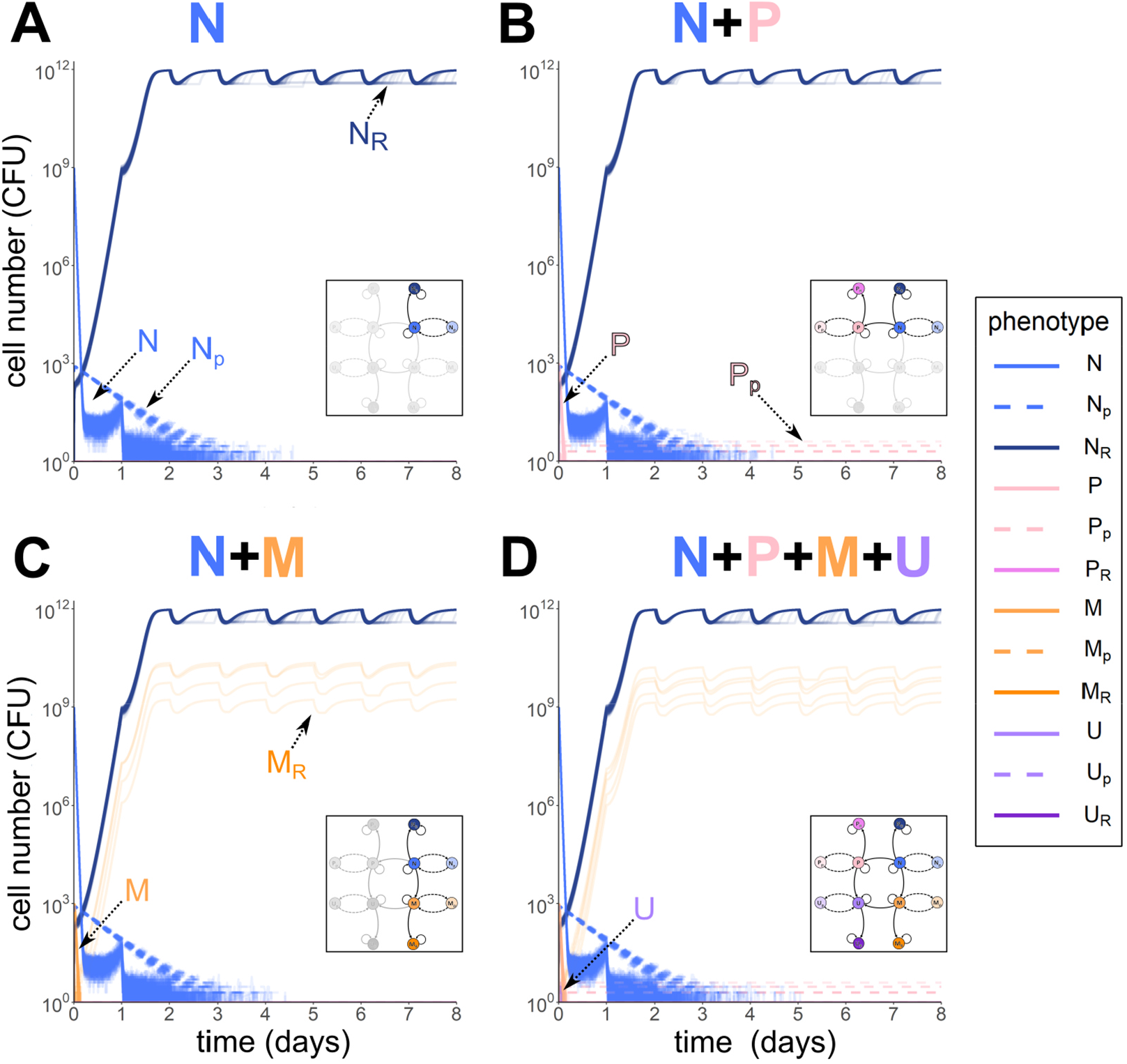
Submodels of resistant, persistent and hyper-mutator phenotypes. Bacterial numbers (CFU) of various phenotypes obtained by stochastic simulations over an 8-day AB treatment with AB administered at 10xMIC (= MIC_R_) every 24 hours. Each figure shows 100 individual simulation runs. **A-C**) display population dynamics of sub-models (as highlighted in the inset). Persister subpopulations are shown as dashed lines, colours of all populations as in Figure 2. **D**) shows the dynamics of the full model shown in Figure 2.

In contrast to our persister simulations (Figure 3A), treatment failure due to persistence is prevalent in chronic infections, even in the absence of resistance, but is often linked to the emergence of high-persisters (P) (Lafleur et al., 2010; Mulcahy et al., 2010; Van den Bergh et al., 2016; Bartell et al., 2020). Compared to N, P are mutants which have a higher rate of switching to persistence (*h*_*F*_-fold) and a lower rate of switching back (*h*_*B*_-fold <<1), resulting in larger persister fractions with a longer ‘life-span’. And indeed, if N can acquire a high-persistence mutation, the emerging P genotype (P+P_p_) is present at treatment failure at low numbers (Figure 3B). Specifically, our simulations show that P rapidly emerges from N, reaches ∼10^3^ CFU, but then rapidly gets killed by the AB. However, in about half of the cases, a very small fraction of P enters the persister state (P_p_) before eradication and persists through the whole AB treatment due to the low back-switching rate from P_p_ (Figure 3B). Note that resistance could emerge from P (P_R_), but in most cases P is eradicated before that happens.

High-persisters are not the only problematic mutants prevalent in chronic infections (Mulcahy et al., 2010). There are also hyper-mutators (M) (Figure 1B), mutants which have an increased mutation rate, and are associated with facilitated evolution of resistance. However, higher mutation rates come at a growth cost (*c*_*M*_), due to the accumulation of detrimental mutations (Montanari et al., 2007). Like with P, we can observe the emergence of the M genotype (M+M_p_) from N at the beginning of treatment at low frequencies (∼10^3^), but it rapidly gets eradicated (Figure 3C). In rare cases M_R_ evolves before M is fully eradicated (∼4%). Although M_R_ and N_R_ are both able to grow under the applied AB dose, due to M_R_ emerging at a later point in the treatment and growing slower than N_R_ by the cost *c*_*M*_, M_R_ only reaches population sizes of around 10^9^ CFU, whereas N_R_ reaches capacity (10^12^ CFU).

Overall, we find that the emergence of the N_R_ genotype is the predominant cause of treatment failure in this regimen, but that the P or M_R_ genotypes can also survive treatment in ∼43% and ∼4% of the cases, respectively. This shows, that both P and M genotypes can confer fitness advantages to N cells and complicate treatment. These results are in line with hyper-mutator frequencies in acute infections, which we compiled from literature (Figure 1B, Text S1), but there is – to our knowledge – no data on high-persister frequencies for acute infections.

Mutations conferring M and P phenotypes do not have to occur independently. As hyper-mutation is by itself detrimental, but is known to facilitate beneficial mutations other than AB resistance (Oliver & Mena, 2010), they could acquire the beneficial high-persistence mutation, which would lead to a higher frequency of the combined genotype than that of hyper-mutators alone. Consequently, we allow for the emergence of a combined mutator-persister genotype (U+U_p_), which has both an increased mutation rate as well as high-persistence switching rates and can evolve from P or M populations through mutation (Figure 2). Further, U can acquire a resistance mutation leading to U_R_. Interestingly, the population dynamics of the full model (Figure 3D) reflect the results of the partial models described above: M and P come up early in the treatment but get quickly eradicated by the AB. U also emerges at the beginning of treatment but reaches even lower population sizes than M or P and is quickly wiped out. The only viable cells left at the end of the 8-day treatment period belong mainly to N_R_, with low numbers of M_R_ and P_p_ populations surviving as well.

### Resistance and high-persistence cause treatment failure at distinct AB concentrations

So far, we only considered one AB concentration (10x MIC) which corresponds to the MIC of the R populations (MIC_R_), but the fitness of the various subpopulations depends on the strength of the selection pressure due to AB. Hence, we investigated which mutant genotype is expected to emerge and establish itself under different AB concentrations. To examine this, we used the full model to determine the probability of a genotype to survive 8 days of treatment for a range of AB concentrations (0-50xMIC). Our simulations show that for sub-MIC survival of all genotypes is possible but starts declining at different concentrations >MIC (Figure 4A, Figure S2). The first genotype to reach zero probability of survival is M, reflecting that hypermutability is costly and without immediate benefit. Then the probability for N survival drops to zero, followed by that of the resistant genotypes P_R_ and U_R_, which already declines for AB concentrations >0.5xMIC_R_, whereas M_R_ survival only reaches zero at about twice the MIC_R_. These differences between resistant genotype survival show that resistance evolution is limited by emergence from the source population (i.e. P, U, M) and that hypermutability can ameliorate this, if only few mutations are necessary. For AB concentrations below ∼2xMIC_R_ survival of N_R_ is almost 100% but drops steeply for concentrations higher than 20xMIC_R_ where N_R_ is replaced as the dominant genotype by the high-persister population (P), which stabilises at around 35% survival for up to 50xMIC. Other subpopulations than N_R_ and P can only be found very rarely at the end of treatment, with e.g. M_R_ surviving in ∼2% of the treatment simulations below MIC and U in <1% in ranges where P populations dominate.

**Figure 4.**
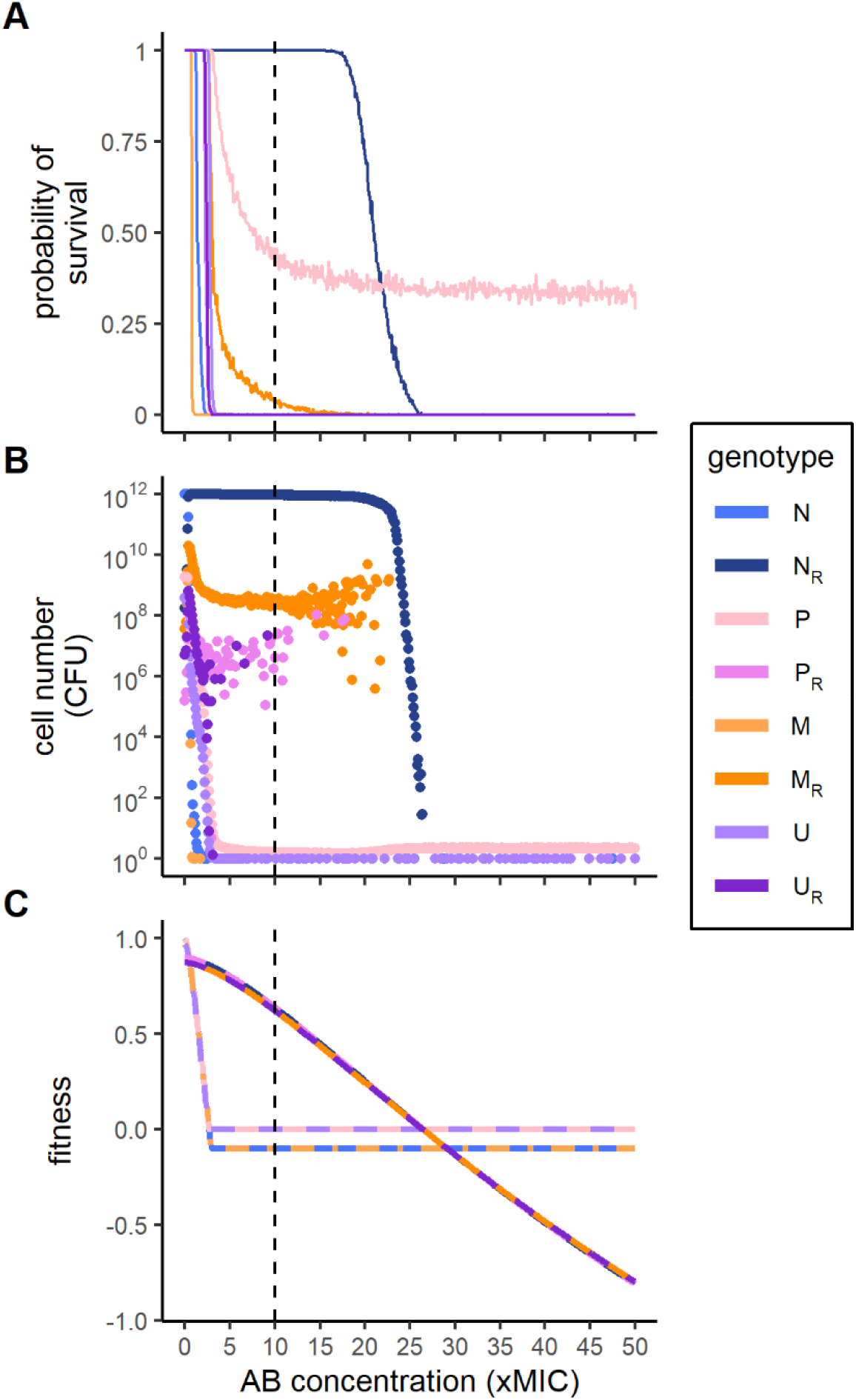
Treatment failure due to wildtype or mutant genotypes. Effect of AB concentration on **A**) the probability of genotype survival at the end of treatment over 1000 simulations runs (see Figure S2 for corresponding probability of treatment failure overall), **B**) the mean absolute cell numbers (CFU) of surviving genotypes, and **C**) deterministic fitness values of the genotypes. **A**) and **B**) summarize the end points of stochastic simulation runs for 8-day AB treatments at various concentrations (0-50xMIC), starting from a population of WT cells (N+N_p_). Fitness of non-resistant populations in **C**) reflects a combination of the underlying growing and non-growing (i.e. persister) subpopulations. Note that in **C**) the fitness curve for N_R_ (P_R_) largely overlaps with the fitness curve for M_R_ (U_R_) for AB concentrations >10xMIC. The vertical dashed line shows MIC_R_.

### In contrast to resistance, persistence causes treatment failure with low numbers of surviving cells

The survival probability does not reflect the absolute pathogen load (mean number of surviving cells) of a certain genotype. For subinhibitory AB concentrations <0.5xMIC, the subpopulations surviving at substantial absolute numbers are diverse (P_R_ = 10^5^-10^8^, M_R_ = 10^7^-10^10^, U_R_ = 10^6^-10^8^ CFU, Figure S3). When N_R_ populations dominate survival, they generally reach carrying capacity (10^12^ CFU). Though differing by orders of magnitude from that, the other resistant genotypes also appear at substantial numbers for AB concentrations up to MIC_R_ (P_R_ and U_R_ at 10^6^-10^7^ CFU) or up to 2xMIC_R_ (M_R_ at 10^9^ CFU) (Figure 4B). In contrast, when P is the dominating population, the total population size drops drastically to less than 10 cells (Figure 4B). These tiny populations largely consist of P and U cells (Figure 4B), and while U survives considerably less frequently than P (Figure 4A), in the cases where U survives, its cell numbers are similar to P. Overall, treatment failure probability above MIC is dominated by N_R_, with bacterial cells reaching carrying capacity, until AB doses exceed MIC_R_ substantially and only persistent, non-growing cells survive at very low numbers. These results do not change if resistant cells are allowed to switch into a persister state as well (Figure S4).

### Differential genotype fitness during treatment is explained by mutational costs and persistence switching rates

To formally understand the change from N_R_ to P as the dominant genotype at high AB concentrations, as well as the low probability of survival of other genotypes, we determine approximate fitness measures for all genotypes as the net growth rate far from carrying capacity (Methods). Since persisters do not grow and arise from a phenotypic – not a genetic – state change, we consider the growing and the non-growing state of a genotype together to calculate its fitness (Text S3). For AB concentrations <0.5xMIC we find the highest fitness for the susceptible genotypes N and P (Figure 4C), as the fitness of all other genotypes is reduced by mutational costs, which outweigh the mutational benefits. However, for AB concentrations between 0.5xMIC and 2.5xMIC_R_ the resistant genotypes N_R_ and P_R_ display the highest fitness, while M_R_ and U_R_ grow slightly slower, due to the cost of hypermutability. Notably, fitness of all genotypes declines with increasing AB concentration, but more slowly for resistant ones. From ∼3xMIC onwards the rate of switching back from persistence determines the fitness of non-resistant genotypes (and hence remains constant at a negative value), with P and U having higher fitness than N and M, due to their lower back-switching rates. In contrast, the resistant genotypes continue to decline with increasing AB concentration until P and U have the highest fitness (which is equal to − *h*_*B*_ * *s*_*B*_ = −10^−4^) for AB concentrations >2.5xMIC_R_. These findings agree with the stochastic simulations of genotype survival and abundance (Figure 4A,C), and arise from the costs of specific mutations as well as the switching rates of persister phenotypes. The residual discrepancy is explained by differences in mutation rates and in the number of mutations necessary for genotypes to emerge, i.e. how fast a genotype can emerge from N (N→M→M_R_ vs. N→M/P→U→U_R_, etc.; Figure 2).

### Mutator-persisters arise from hyper-mutators during acute infections and from high-persisters at relapse

Even though with very low probability and numbers, cells combining hyper-mutation and high-persistence phenotypes (U) can survive at the end of the treatment. This could result in subsequent treatments being ineffective due to 1) the larger fraction of persistent subpopulations, or 2) the higher mutation rates facilitating the emergence of problematic mutations like resistance, or 3) a combination of both. Hence, the emergence of U deserves a closer inspection, specifically, which population they originate from, i.e. do they emerge from the fitter P cells or do M cells acquire the beneficial high-persistence mutation? By tracking the mean cumulative mutation events of either M or P populations producing U cells over the course of treatment, we find that, surprisingly, despite the high survival rate of P, contributions from M to U are higher than the contributions from P for all AB concentrations (Figure S5). As the cumulative contributions to U do not necessarily reflect the establishment of the resulting U population (Text S4), we ran simulations where either only M or only P could mutate to U. We found that the U population only emerges in a similar manner as before (Figure 4B), if M-to-U mutations are allowed (Figure S6), which makes M cells the main source population for U over the course of 8-day treatments, i.e. during acute infections.

However, P subpopulations are likely to become problematic after AB treatment subsides, as they provide a pool of viable cells, which can regrow in the absence of ABs and cause relapse of the infection (Lewis, 2010). To capture this, we model the regrowth of populations surviving 8-day AB treatment at less than 10^5^ CFU, which only occurs at AB concentrations higher than 1.5xMIC_R_ (Figure 4). Here, we assume that >10^5^ cells are clinically detectable and would result in treatment continuation (Figure 5A). More than 90% of the ‘non-detectable’ surviving populations can regrow to at least 10^6^ CFU within 10 years, causing a relapse of infection (Figure 5B). The cell numbers of different genotypes at relapse, mainly correspond to P and U, as well as small P_R_ and U_R_ populations (Figure 5C), suggesting that P and U subpopulations increasingly play a role in recurring infections as indicated by clinical studies (Mulcahy et al., 2010; Bartell et al., 2020). Notably, the N_R_ genotype only causes relapse at relatively low frequencies and for a very narrow range of AB concentrations (Figure 5C). These populations however result in relapse within days (Figure S7). This is in stark contrast to the time until relapse caused by P and U, which are in a persister state at the end of treatment and must first switch back to a growing state – which on average happens within a year (median = 37 weeks; Figure S7). When considering acute treatment and relapse combined, the total contributions of P to U are higher than those from M (Figure S5) as M does not survive acute treatment at AB concentrations relevant for relapse simulations (Figure 4A). Hence, while M is the main source population of U during acute infection treatment, after AB treatment ends, P becomes the main source population.

**Figure 5.**
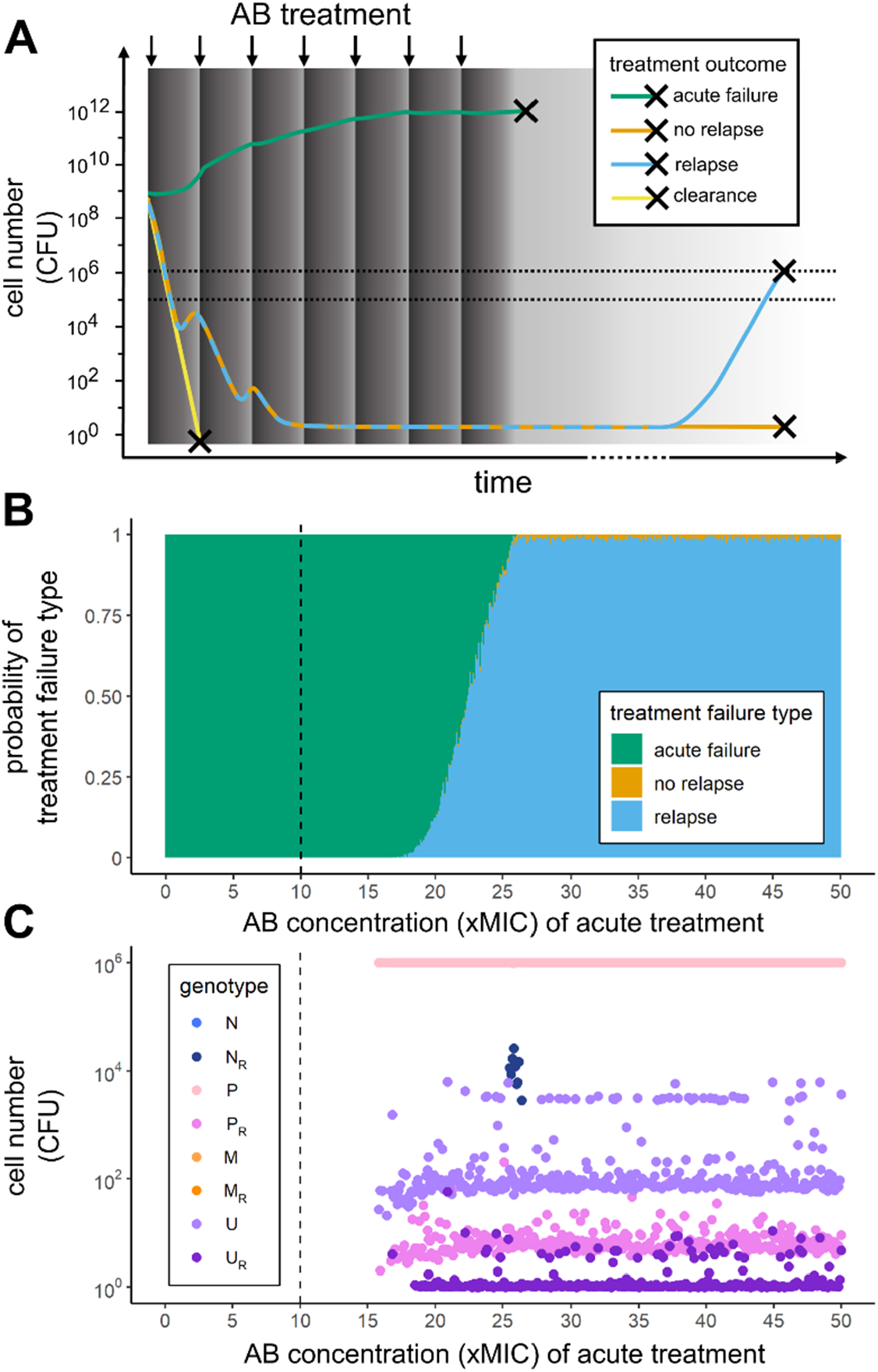
Relapse is mainly caused by persister phenotypes. **A**) Bacteria can either get cleared (yellow line) or survive 8-day AB treatment (dark grey area; periodic dosing is indicated by arrows and AB decay as a gradient). Surviving bacteria either cause immediate, acute treatment failure (>10^5^ CFU, lower dashed line) at the end of treatment (green line) or, due to regrowth from <10^5^ CFU, which leads either to relapse (>10^6^ CFU as indicated by the upper dashed line; blue) or no relapse (remaining below 10^6^ CFU; orange) over the course of 10 years. **B**) Probability of each of the three treatment failure types to occur for a range of AB concentrations used in the treatment (0 – 50xMIC). **C**) Mean cell numbers (CFU) of each genotype at relapse. Vertical dashed lines in **B**) and **C**) show MIC_R_.

## Discussion

Acute and especially chronic infections with bacterial pathogens like *P. aeruginosa* or *E. coli* show a concerning frequency of mutator and high-persistence phenotypes, both of which are known to facilitate the evolution of antibiotic resistance (Oliver & Mena, 2010; Levin-Reisman et al., 2017; Rodriguez-Rojas et al., 2021). Further, given their high prevalence in chronic infections (Prunier et al., 2003; Labat et al., 2005; Lafleur et al., 2010; Mulcahy et al., 2010; Schumacher et al., 2015; Bartell et al., 2020), these two phenotypes could possibly combine and aggravate the problem. In this study, we used a stochastic population model to disentangle the complicated emergence and interplay of phenotypic and genotypic survival strategies under AB treatment by investigating the evolutionary dynamics of hyper-mutator (M), high-persister (P) and resistance (R) genotypes over the course of AB treatment at various concentrations and during relapse after treatment ends.

We find that for relatively low (but higher than MIC) AB concentrations treatment failure is certain and caused by R genotypes which grow to carrying capacity by the end of the treatment (Figure 4A,B). In contrast, for antibiotic concentrations much higher than MIC_R_ treatment failure happens only in about a third of the cases and is caused by high-persistence genotypes in persister state at very low population sizes. This behaviour is explained by our deterministic fitness calculations, which show that for AB concentrations >2.5xMIC_R_ fitness of the high-persistence genotypes, P and U, is larger than that of the resistant genotypes as persisters are unaffected by high AB concentrations (Figure 4C). P and U as the main cause for treatment failure at high AB concentrations fits with the clinically observed “paradox of chronic infections” in the absence of resistance (Lewis, 2010; Mulcahy et al., 2010), especially as peak AB concentrations reached in the treatment of cystic fibrosis (CF) patients with inhaled, nebulized ABs are high and on average >2.5xMIC_R_ (Eisenberg et al., 1997). Our findings suggest that treatment failure via high-persistence is most likely to occur if sufficient resistance cannot easily evolve. This means that AB concentrations targeted at exceeding MIC_R_ might instead select for ‘hidden’ chronic infections due to persisters. Hence, persisters could specifically be targeted at later time points of the treatment with anti-persister drugs (Defraine et al., 2018). This strategy could be tested under laboratory condition by treating bacterial populations with high AB doses followed by anti-persister drugs and comparing it to potential regrowth from cultures without anti-persister treatment.

The total population size of P and U during AB treatment can only decline if the persisters switch back to the growing state, which for high-persistence mutants occurs very slowly (Figure S7), hence, they die more slowly than other genotypes. Accordingly, we see that persister subpopulations primarily prolong the time to clearance and constitute a substantial fraction of the bacterial population only at later stages of treatment when all (or most) non-persisters have died (Figure 3). The wildtype switching rates used here (Balaban et al., 2004) might even underestimate persister survival as they lead to shorter lifetimes of persister subpopulations than found by Svenningsen et al. (2021), who reported that *E. coli* persisters can survive at least 7 days of antibiotic exposure. Using random parameter sampling and linear discriminant analysis (LDA) to investigate the influence of transition rates between genotypic and phenotypic subpopulations (i.e. mutations and switching rates; Methods), we find that wildtype (N) switching rates to (s_F_) and from (s_B_) persistence influence treatment outcome substantially (Figure S8). Particularly, s_B_ has a large impact, with higher back-switching rates leading to more clearance, which fits well with the role of s_B_ in determining the fitness of high-persisters (Figure 4C). This should be even more important in clinical infections, where multiple stressors are present and bacterial doubling times are generally much slower than under laboratory conditions. Yet, empirical determination of back-switching rates from persistence under different conditions and for different genotypes is scarce so far. Further, the commonly used time frames of 8-24h might not be sufficient to empirically investigate persistence - and especially high-persistence - dynamics appropriately.

Persistence and high-persistence have been shown to facilitate the evolution of resistance over the course of antibiotic exposure under laboratory conditions (Levin-Reisman et al., 2017; Rodriguez-Rojas et al., 2021). Interestingly, in our simulations P does not facilitate the evolution of resistance during the treatment of acute infections as illustrated by the low probability of survival of the P_R_ genotype at the end of treatment (Figure 4AB). This is reflected in our LDA, where an extremely low percentage of simulations with random parameters result in P_R_ as the dominant genotype (Figure S8A). Instead, P enables survival at AB concentrations where antibiotic resistance is not viable anymore in our regimen (Figure 4C). We find that this reasoning is robust for a wide range of parameter sets (Figure S8D) as higher mutation rates to resistance (µ_R_) and higher AB concentrations (A_max_) have the largest influence on pushing treatment outcome towards failure due to resistance or failure due to persistence respectively. Empirical evidence for the distinction between persistence and resistance in causing treatment failure comes from chronic *P. aeruginosa* infection in CF patients, where high-persistence phenotypes were prevalent, but only some were additionally resistant (Mulcahy et al., 2010). This potentially indicates that resistance via chromosomal mutations, as simulated here, might be less easily attainable or less beneficial in disease settings than in the lab. Additionally, the discrepancy between clinical findings and laboratory experiments regarding resistance-facilitation by persisters can partially be explained by experimental limitations: Directed evolution experiments (Levin-Reisman et al., 2017) use only comparatively low AB concentrations for relatively short time frames while simultaneously allowing for long AB free regrowth periods. Accordingly, while we do not find that persistence facilitates resistance evolution over the course of acute treatment, when we consider relapse after AB treatment ends, we indeed see P_R_ cells coming up (Figure 5B). However, our assumption that persisters cannot mutate is likely over-simplistic and currently remains an open question in the field, but there are empirical studies indicating that persisters can still be metabolically active (reviewed in Kim & Wood, 2016) and might even increase mutation rates (Windels et al., 2019). Further, we are not considering plasmid-borne resistance in this study, which could speed up resistance emergence, but it is unclear if plasmid conjugation occurs in persister subpopulations.

Our simulations show that, in comparison to P, M is much more likely to facilitate the evolution of resistance (Figure 4), which agrees with theoretical (Travis & Travis, 2002) and experimental studies (Giraud et al., 2002). However, while M_R_ readily evolves, it emerges later than N_R_ (Figure 3C) and grows at a slower rate due to fitness costs of hyper-mutation, which results in M_R_ reaching lower population sizes than N_R_. Hence, if resistance evolves readily enough from N, hyper-mutators cannot dominate the population due to resistance-facilitation. However, it is still possible that M could acquire other beneficial mutations mitigating its cost, which are not accounted for in our model (Oliver & Mena, 2010). Further, in clinical conditions M subpopulations might already be present at the onset of AB treatment, speeding up the emergence of M_R_. Nonetheless, our simulation results for M survival of acute treatment (Figure 3, Figure 4A) are in line with hyper-mutator frequencies found in acute infections, but significantly lower than those found in chronic infections, indicating a potential role of acquiring beneficial non-resistance mutations (Figure 1B).

In addition to the individual impact of high-persistence and hyper-mutation on the potential for treatment failure, we investigated the emergence of a combined genotype (U). Generally, the dynamics of the full model (Figure 3D) mirror the dynamics of the individual sub-models for M and P regarding their survival probability and end population size (Figure 3B,C). This is *a priori* not obvious for such a complicated system involving various genotypes and phenotypes and gives hope that studies of isolated systems can provide information about more complex combinations. Further, we find that U cells mainly emerge from the M population (Figure 4A,B, S6). Hence, in accordance with empirical studies (Oliver & Mena, 2010), we find that M could hitch-hike a beneficial mutation, here the high-persistence mutation, to offset the cost of hyper-mutation. Interestingly, since the adaptive value of high-persistence comes from the non-growing persister state, the growth cost of hyper-mutation could matter less in U. Thus, M could facilitate rapid emergence of U early during treatment, which would allow the hyper-mutation to get fixed at minimal cost via U – as opposed to the situation where M acquires R and enters growth competition with other R genotypes (Figure 3C).

Notably, we find that the subpopulation dynamics change between treatment failure of acute infection and relapse after treatment is discontinued. While U is most likely to arise from M during the treatment of acute infections, we find that during regrowth of small surviving populations, more U emerge from P than from M (Figure S5). Therefore, high-persistence mutants, such as *HipQ* (Wolfson et al., 1990; Balaban et al., 2004), might not facilitate evolution on a short time scale, but rather on a longer time scale after stress subsides. Since both hyper-mutators and high-persisters are generally associated with chronic infections it is crucial to consider their dynamics not only during, but also following, treatment, i.e. during potential relapse (Figure 5). Specifically, high-persisters have been proposed to cause recurring infections by providing a small pool of surviving cells, which start to regrow once AB concentrations subside (Lewis, 2010). This is in agreement with our findings, where relapse from small surviving populations is common (Figure 5A) and predominantly caused by P (Figure 5B). Additionally, we find P_R_, U and U_R_ genotypes in small numbers (Figure 5B), showing that P does not only survive high AB concentrations to cause relapse, but also facilitates the emergence of resistance in the absence of ABs, as has been shown experimentally (Levin-Reisman et al., 2017).

Our relapse model likely overestimates the time until persisters wake up in the absence of ABs, and therefore the time until relapse (Figure S7), as we assume a constant (and especially for hyper-persisters very slow) back-switching rate. This assumption corresponds to so-called ‘spontaneous persistence’ but neglects ‘triggered persistence’ (Balaban et al., 2004; Balaban et al., 2019), which is characterized by switching rates that are dependent on “trigger” stressors, such as starvation (Svenningsen et al., 2021) or ABs (Dörr et al., 2009). Therefore, considering triggered persistence could lead to faster switching back from persistence in the absence of ABs. Disentangling the effect and magnitude of multiple stressors on triggered switching is complicated and parameterization attempts suffer from danger of overfitting (Van den Bergh et al., 2016; Carvalho et al., 2017), which is likely the reason why – to our knowledge – no parameter estimates for triggered switching are available for high-persisters. Overall, our model simulations provide a conservative estimate of the probability of relapse, which could be higher with triggered persistence.

Lastly, persister frequencies show a high amount of variation in empirical studies, even in the presence of the same AB (Figure 1D), which could be caused by stochasticity or differences in cellular physiology and persistence-causing mechanisms (Allison et al., 2011; Kint et al., 2012). Notably, when grouped by mechanism of action, antimicrobials which target the bacterial membrane display the lowest persister frequencies (Salcedo-Sora & Kell, 2020). All membrane-targeting antimicrobials analysed by Salcedo-Sora and Kell (2020) were antimicrobial peptides (AMPs), which might indicate reduced persister formation or survival with AMPs as compared to ABs. Running our simulations with AMP-like pharmacodynamic parameters (Methods, Table S1), we find drastically lower survival of high-persistent and resistant bacteria (Figure S9) than for AB-like pharmacodynamics (Figure 4). This is due to AMPs killing bacteria faster than ABs, allowing less opportunity for mutation emergence and switching into the persister state. This suggests that AMPs can decrease the chance of high-persister (and mutator-persister) emergence, and thereby the probability of relapse, while at the same time allowing for less resistance evolution (Yu et al., 2018).

In conclusion, we find that high-persistence and hyper-mutant genotypes mainly act independently and on different timescales with hyper-mutator cells facilitating the emergence of resistance over the course of 8-day AB treatment, and high-persistence enabling survival at high AB concentrations and resistance evolution after treatment ends. Accordingly, we find that the emergence of the combined mutator-persister genotype is driven by different populations during acute treatment (M) and during relapse (P). Generally, the treatment AB dose relative to the MIC of the resistant population is an important determinant for the selection of different genotypes: while genetic resistance leads to immediate treatment failure for AB levels up to 2.5xMIC_R_, high-persistence dominates at higher AB levels, leading to relapse after drug removal. Hence, particularly the interplay of genotypes and phenotypes needs to be studied in environments with fluctuating stressors. More broadly, our modelling framework is not limited to AB treatment of bacterial infections but can be applied to other diseases, where drug efficacy is inhibited by genotypic and phenotypic mechanisms, such as in fungal infections (Lafleur et al., 2010; Healey et al., 2016) or cancer (Sharma et al., 2010; Campbell et al., 2017). Our results suggest that treatment strategies should consider the different timescales at which various AB-escape mechanisms operate to reduce the risk of treatment failure or relapse.

## Methods

### Stochastic population model

We investigate the relative importance of high-persistence, hyper-mutation and resistance mutations (leading to P, M and R subpopulations, respectively) – and all their combinations – over the course and after discontinuation of AB treatment by using a stochastic population model (Figure 2). Our model incorporates pharmacokinetic and pharmacodynamic functions to realistically simulate AB treatment (Eq. 1) and the distinct effects of ABs on our respective subpopulations. As proposed by Balaban et al. (2004) we describe phenotypic persistence as a two-subpopulation process. For each of our AB-susceptible genotypes (the wildtype N, M, P and mutator-persisters U) we model a growing subpopulation, which is affected by ABs (Eq. 2), and a non-growing (i.e. *ψ*_*max*_ = *0*) persister subpopulation, that is unaffected by ABs. The transitions between growing and persister subpopulation happen stochastically at rates *s*_*F*_ (N → N) and *s*_*B*_ (N → N) for N and M, or *h*_*F*_ *\* s*_*F*_ (P → P_*p*_) and *h*_*B*_ *\* s*_*B*_ (P → P_*p*_) for P and U. In our model switching rates are constant (i.e. not environment-dependent) and we assume that resistant subpopulations (N_R_, P_R_, M_R_ and U_R_) do not generate persisters, as we found that persistence does not convey any additional benefit to already resistant bacteria (Figure S4).

For our starting conditions we assume that only the susceptible WT genotype, consisting of its two phenotypic states, N and N_p_, is present at 1×10^9^ colony forming units (CFU) total (N+N_p_), with N and N_p_ being in equilibrium, according to their respective switching rates. The ratio 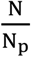 at equilibrium is obtained by determining the dominant eigenvector of the analytical solution of the two-population persister model from Balaban et al. (2004) in the absence of ABs and with our respective parameters (see also Rodriguez-Rojas et al., 2021).

Growth of the non-persister populations is limited by the overall carrying capacity *K*, which, together with a constant natural death rate *d*, results in realistic competition between genotypes. Since our mutations are coupled to growth, the death rate also enables mutations to occur after capacity is reached. P and M mutants can only arise by mutation from the susceptible growing N population at rates µ_*P*_ and µ_*M*_ respectively, and U from the susceptible growing P and M populations at rates µ_*P*_ and µ_M_ respectively. To investigate the relative contribution of M and P to the emergence of U, we separately quantify mutation events from M and P leading to U when mutation to U is only possible from either M or P.

Resistant mutants N_R_ and P_R_ emerge by mutation from N and P at mutation rate µ_*R*_, whereas M_R_ and U_R_ have elevated mutation rates and arise from M and U at *h*_µ_ *\** µ_*R*_. The maximal growth rate of hyper-mutator populations (M, M_R_, U and U_R_) is reduced by the cost of hyper-mutation (*C*_*M*_): 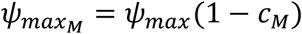 (Text S1). Similarly, the growth of resistant populations is reduced by the cost or resistance (*C*_*R*_): 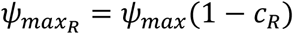. Note that for M_R_ and U_R_ these costs are multiplicative. See Text S2 for the corresponding system of ordinary differential equations. We calculate the probability of genotype survival at the end of the treatment as the fraction of simulation runs per AB concentration where the genotype population size is larger than zero.

### Pharmacokinetic and Pharmacodynamic functions

We model bactericidal AB treatment with periodic dosing intervals and exponential AB decay, as shown in Figure S1, by using the pharmacokinetic function

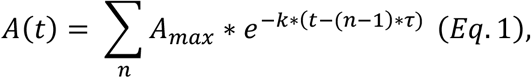

with 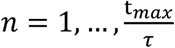 representing the number of dosing events. For our simulations we model 8-days of daily AB treatment (*t*_*max*_ = 192h, *r* = 24h) and examine a broad range of drug concentrations *A*_*max*_ covering 0xMIC to 50xMIC of the susceptible populations. Note that due to drug decay, the concentration *A*_*max*_ denotes the peak AB concentration, but will be generally referred to as drug concentration in the main text. The decay parameter *k* is fixed for all our simulations, except stated otherwise (Table S1).

The effect of an AB on a bacterial population is defined by the pharmacodynamic function:

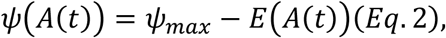

with 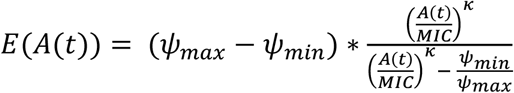 (*Eq. 3*) (Zhi et al., 1986; Regoes et al., 2004).

The parameters *ψ*_*max*_ and *ψ*_*min*_ describe the maximal and minimal net growth rates in the absence (*ψ*_*max*_ = *ψ*(*A* = *0*)) or the presence of high amounts of ABs (*ψ*_*min*_ = *ψ*(*A* → *∞*)). At AB concentrations equal to their minimal inhibitory concentration (MIC) bacterial populations do not grow (*ψ*(*A* = *MIC*) = 0). Resistant populations are assumed to have MIC_R_ = 10xMIC, meaning that their growth stops at a 10-fold higher AB concentration than for susceptibles. The Hill parameter *K* determines the steepness of the pharmacodynamic curve described by Eq. 2, which reflects the sensitivity of bacterial growth to AB concentration changes. See Table S1 for all parameters values of AB simulation treatments as well as for Antimicrobial Peptide (AMP) pharmacodynamics, where the latter are characterized by lower *ψ*_*min*_ and steeper *K*.

### Deterministic Fitness Measures

To investigate which populations should be fittest for different values of A_max_, we separately calculate long-term growth rates for each genetically unique population as approximate fitness measures. For all resistant populations (N_R_, M_R_, P_R_ and U_R_) this is achieved by integrating the pharmacodynamic functions (Eq. 2), which describe the net growth rate plus the natural death rate d (Eq. S6, S9, S12), over one treatment period (*0* to *τ*) and dividing by *τ* to derive the mean growth rate per hour (see Text S3 for closed integrals). Note that we assume that population sizes are far from the carrying capacity, which allows us to neglect part of the logistic growth.

For the fitness calculations of genotypes with persister phenotypes we need to consider both sub-populations together as they are linked via constant switching and, especially under AB treatment, both contribute to the fitness of the genotype. These two phenotypic states are affected by ABs to a different extent, one growing and susceptible, the other one not-growing and not affected by ABs. Hence, we calculate the analytical solutions for the susceptible populations for combined growing/persister-pairs: N+N_p_, P+P_p_, M+M_p_ and U+U_p_. The analytical solution of these ordinary differential equations is *S*(*t*) = N(*t*) *+* N_*p*_(*t*) = *g*_*+*_ * *e*^*λ+t*^ *+ g*_-_ * *e*^*λ−t*^, with 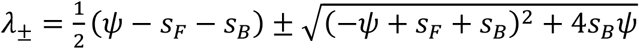 (see Balaban et al. (2004) and Komarova and Wodarz (2007), here adapted to our notation). Average net growth rates *ψ* of the individual pairs are derived via integration of the pharmacodynamics function of the growing subpopulation, including growth costs where appropriate (M, U), as described for resistant populations above (Text S3). The factors *g*_*+*_ and *g*_-_ can be calculated from the initial starting population sizes of N(0) = N_0_ and 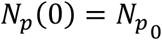 with 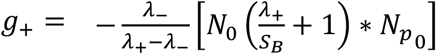 and 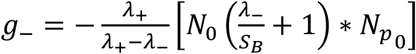. For relevant parameters, that is *s*_*F*_ > 0 and *s*_*B*_ > 0, it follows that *λ*_*+*_ > 0 for *ψ* > 0 (net growth), *λ*_*+*_ < 0 for *ψ* < 0 (net killing), and if *λ*_*+*_ < 0, then |*λ*_*+*_| ≤ |*λ*_-_|, *λ*_-_ < 0 and *g*_*+*_ ≥ 0. From these properties, it follows that the long-term behaviour or net growth rate of the growing/persister-pair is determined by 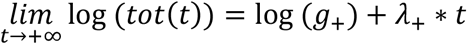, where *t* stands for time and *tot* for the growing/persister-pair (see also Komarova and Wodarz, 2007). The asymptote described by 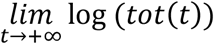 is best understood as the second phase of the biphasic killing curve (Figure 1A), whereby *λ*_*+*_ describes the slope and *log* (*g*_*+*_) the y-intercept. Since we assume small population sizes, the effect of *g*_*+*_ can be neglected and *λ*_*+*_ is the main descriptor for long-term behaviour of growing/persister-pairs.

### Relapse simulations

High-persisters and hyper-mutators are both prevalent in chronic infections and persisters have been proposed to cause relapse of infection even in the absence of resistance (Lewis, 2010). Hence, in addition to the 8-day AB treatment described above, we also simulate how and when surviving bacterial populations can cause an infection to relapse after AB treatment ends. For these relapse simulations we only consider treatment outcomes, where treatment failure is not apparent, which we define by the total surviving population size being <10^5^ CFU. If >10^5^ cells survive the treatment, we consider that as an apparent, acute treatment failure and do not run a relapse simulation. In clinic reality, due to individual differences in patients and infections, determining such a cut-off is much more complicated and as such beyond the scope of this work. However, 10^5^ CFU is in line with clinical detection limits, for example for the diagnosis of urinary tract infections (Schmiemann et al., 2010). The regrowth of these small surviving populations is simulated according to the equations outlined above, but in the absence of AB administration (only considering the decaying, leftover AB from the treatment), until a total bacterial population size of 10^6^ CFU is reached, or alternatively for a maximum time of 10 years. We assume that relapse will only be noticeable at pathogen loads that are an order of magnitude higher than our detection limit of 10^5^ CFU as monitoring of an ongoing infection likely leads to detection of lower bacterial numbers.

### Parameter sensitivity analysis

To assess how sensitive our simulation results are to our specific choice of parameters, we investigate the effect of the main six parameters of interest regarding transitions between subpopulations: the switching rates of the wild type (*s*_*F*_, *s*_*B*_), the mutation rates to resistance (*µ*_*R*_), to hyper-mutation (*µ*_*M*_) and to high-persistence (*µ*_*P*_), and the AB concentration (*A*_*max*_). Note that *µ*_*M*_ and *µ*_*P*_ were explored in correlation with *µ*_*R*_ by varying the fold-change difference to *µ*_*R*_ (Table S1). We randomly sample each of these six parameters 100,000 times using Latin Hypercube Sampling (LHS) (Carnell, 2020) to ensure efficient and complete coverage of our designated parameter ranges (Table S1). *A*_*max*_ was sampled from a uniform distribution and all other parameters from log-uniform distributions. Briefly, LHS divides the range of each parameter into quantiles equal to the number of samples, here 100,000, and randomly samples once for each quantile. These 100,000 samples of each parameter are then combined randomly into 100,000 parameter sets. For each of these sets we run 100 simulations of 8-day AB treatment (with the other parameters as for the main simulations; see Table S1). To get an overview of the results, we first consider the dominant genotypes at the end of treatment as percentages of the 10,000,000 simulations (Figure S8A). Dominant genotypes were defined as the genotype with the absolute highest cell number and simulations where two genotypes had the highest cell number were labelled ‘equal’. If no bacterial cells survived ‘clearance’. For determining parameter effects, we chose however three broader classes of treatment outcome (Figure S8B), according to our main results in Figure 4: ‘clearance’ (no surviving bacteria), ‘resistance’ (majority of surviving bacteria are resistant genotypes) and ‘persistence’ (majority of surviving bacteria are persistent/susceptible). Here, ‘majority’ is again defined as the highest cell number at the end of treatment for all resistant genotypes or all non-resistant genotypes combined. In 90% of the cases the cell numbers between these two classes differ by orders of magnitude. Note, that ‘persistence’ here includes all susceptible genotypes since all simulated AB concentrations are ≫MIC where susceptibles can only survive in persister state (Figure 4C). To assess the individual effects of our six parameters of interest on treatment outcome we use Linear Discriminant Analysis (LDA) on these classes (Tepekule et al., 2017). Simplified, LDA projects the multi-dimensional data onto a 2D space in a way that maximally separates the individual classes from each other.

### Implementation

All simulations, analysis and plots were done in R version 3.6.0. Stochastic simulations were implemented via the Gillespie algorithm using the R-package *adaptivetau* (Johnson, 2019). To test the accuracy of our simulation results, we used different tolerance levels for the relative rate changes in step size selection, which did not change our results notably. The stochastic simulations were run 1000 times for each AB concentration (0-50xMIC at 0.1 steps) for 8 days for acute treatment and 10 years for relapse. Analytical solutions of the population models were determined by using Matlab version R2020b.

## Data Availability

Data will be made publicly available on github.

## Acknowledgements

We thank J. Baer, S. Lehtinen and J. Rolff for useful discussions and comments on the manuscript. This work was supported by the Swiss National Science Foundation (Grant 310030B_176401) and by an ETH Zurich Postdoctoral Fellowship (19-2-FEL-74) received by CI.

