## Supplemental Material for "Assessing the importance of resistance, persistence and hyper-mutation for antibiotic treatment success with stochastic modelling"

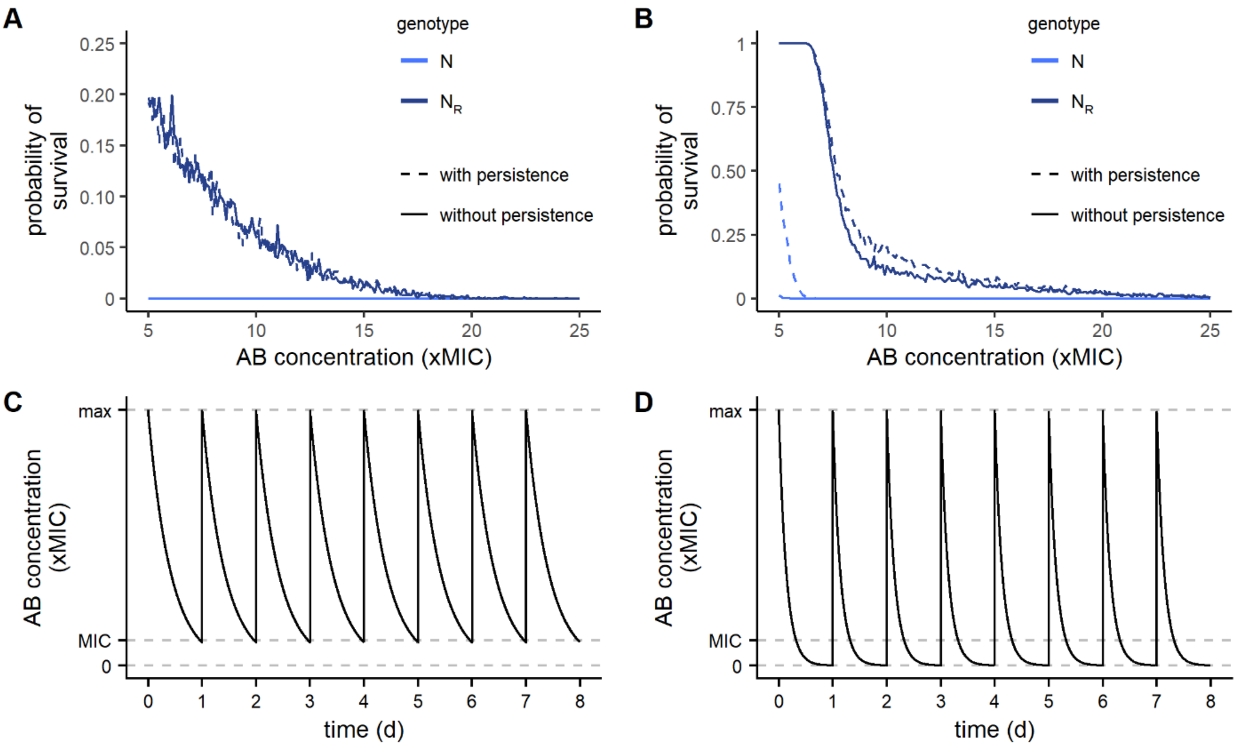


**Figure S1. The effect of persistence on the emergence of resistance**. Probability of genotype survival over the course of 8-day AB treatment is shown for N (light blue) or $N_R$ (dark blue) cells with (dashed line) or without (solid line) persister formation for AB decay rates of **A**) 0.1 or **B**) 0.3 $h^{-1}$. Initial cell densities ($N+N_p$) are $10^6$, which visualizes the effect more clearly than $10^9$ (as used for the main simulations). **C**) and **D**) show the corresponding pharmacokinetics of AB concentration over the treatment period for 0.1 and 0.3 $h^{-1}$ decay, respectively. If AB concentrations fall below the MIC, they allow cells that switch back from $N_p$ to N to survive and enable evolution of resistance.

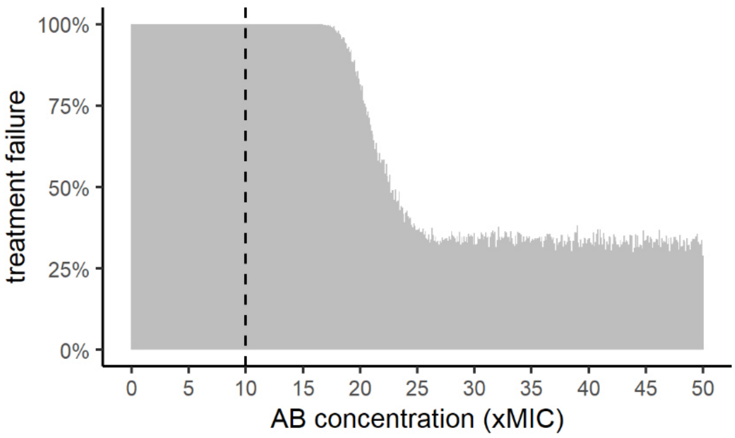


**Figure S2. Probability of treatment failure for different AB concentrations.** Percentage of failed treatments over 1000 simulations with the full model version per AB concentration (0-50xMIC) are shown. Treatment failure is defined as ≥1 viable bacterial cell present at the end of the 8-day AB treatment. The black, dashed line shows $MIC_R$.

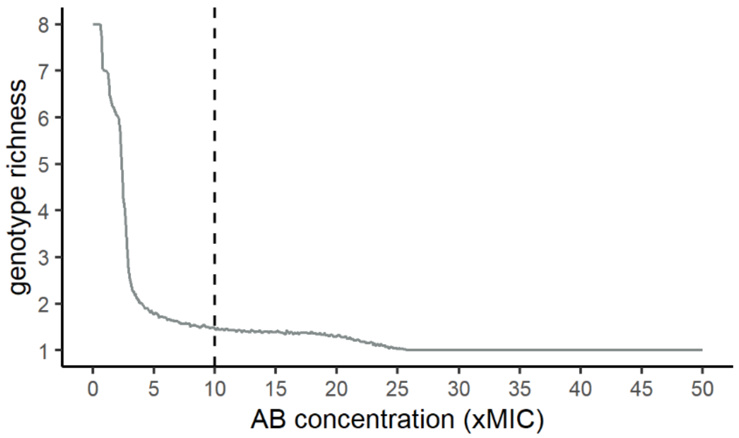


**Figure S3**. **Genotype richness is AB dependent.** Genotype richness is calculated as the mean number of surviving genotypes present at the end of 8-day AB treatment for a range of AB concentrations.

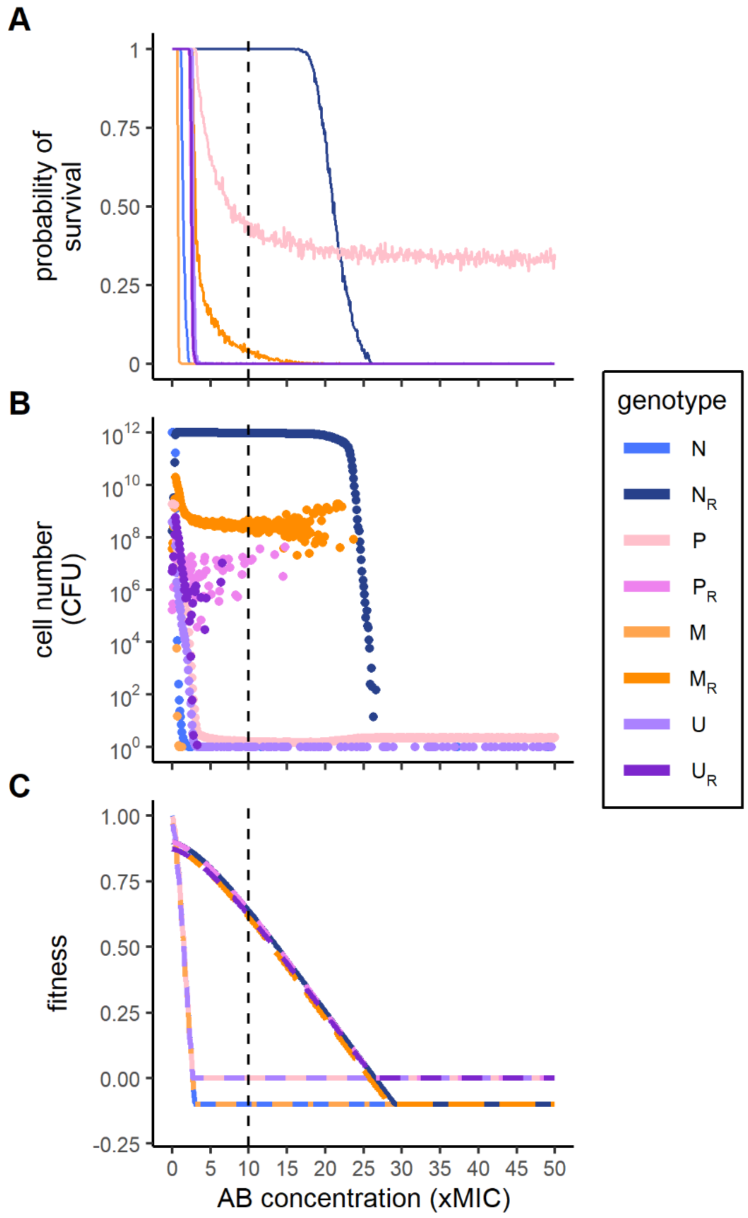


**Figure S4. Treatment simulations including persister subpopulations of resistant mutants**. The ability of resistant genotypes (subscript R) to generate persisters does not change the qualitative results (compare to Figure 4 A,B) of the effect of AB concentration (0-50xMIC) on **A**) the probability of genotype survival at the end of an 8-day AB treatment over 1000 simulations runs, and **B**) the mean absolute cell numbers (CFU) of surviving genotypes. **C**) The deterministic fitness values of the resistant genotypes are however now bounded from below by the persister fitness instead of declining further (Figure 4C). Note that in **C**) the fitness curve for $N_R$ ($P_R$) largely overlaps with that for $M_R$ ($U_R$). This effect does not change stochastic simulation outcomes (A,B) substantially because resistant populations large enough to produce a significant amount of persisters will dominate regardless of persistence. Vertical dashed line shows $MIC_R$.

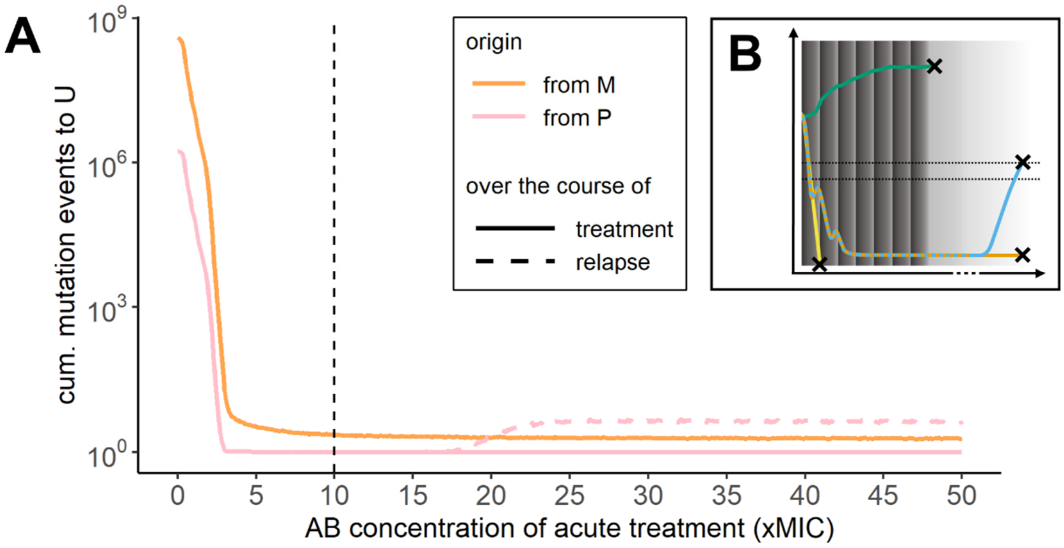


**Figure S5. Mutation events from hyper-mutator (M) and high-persister (P) subpopulations leading to mutator-persisters (U). A**) Shown are cumulative mutation events to U (counts) for a range of AB concentrations (0-50xMIC) as solid lines (orange from M and pink from P populations) if they occur during AB treatment (dark grey area in B) and as dashed lines if they occur during relapse (dark grey + light grey area in B). The black dashed line shows $MIC_R$. **B**) The AB treatment lasts for 8 days during the acute infection (dark grey area) and is followed by an AB-free period (light grey area) during which surviving bacterial cells can grow up to cause relapse (defined as >$10^6$ CFU, upper dashed line; Methods).

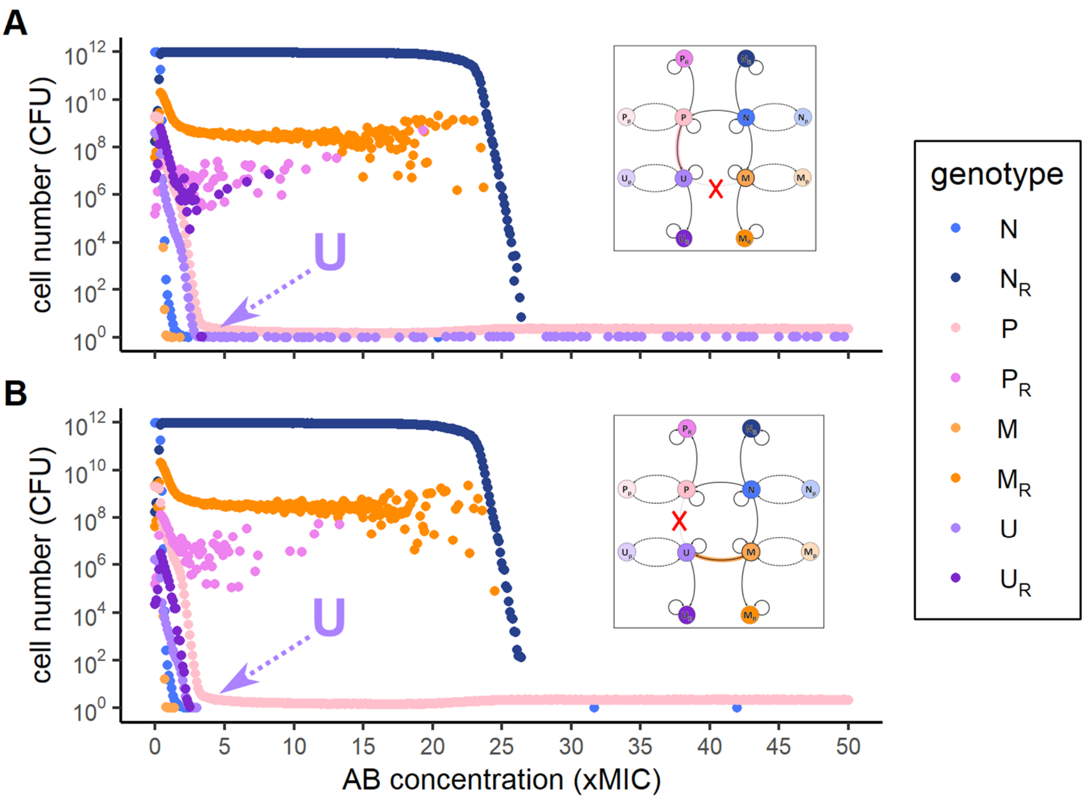


**Figure S6. Mutator-persisters (U) arise mainly from Mutators (M), not from high-persisters (P).** Mean population sizes of genotypes at treatment failure show that if **A**) mutations from P to U are not possible, U cells can still be found for a wide range of AB concentration. In contrast, if **B**) mutations from M to U are not possible, U cells are diminished for low and lost for high AB concentrations.

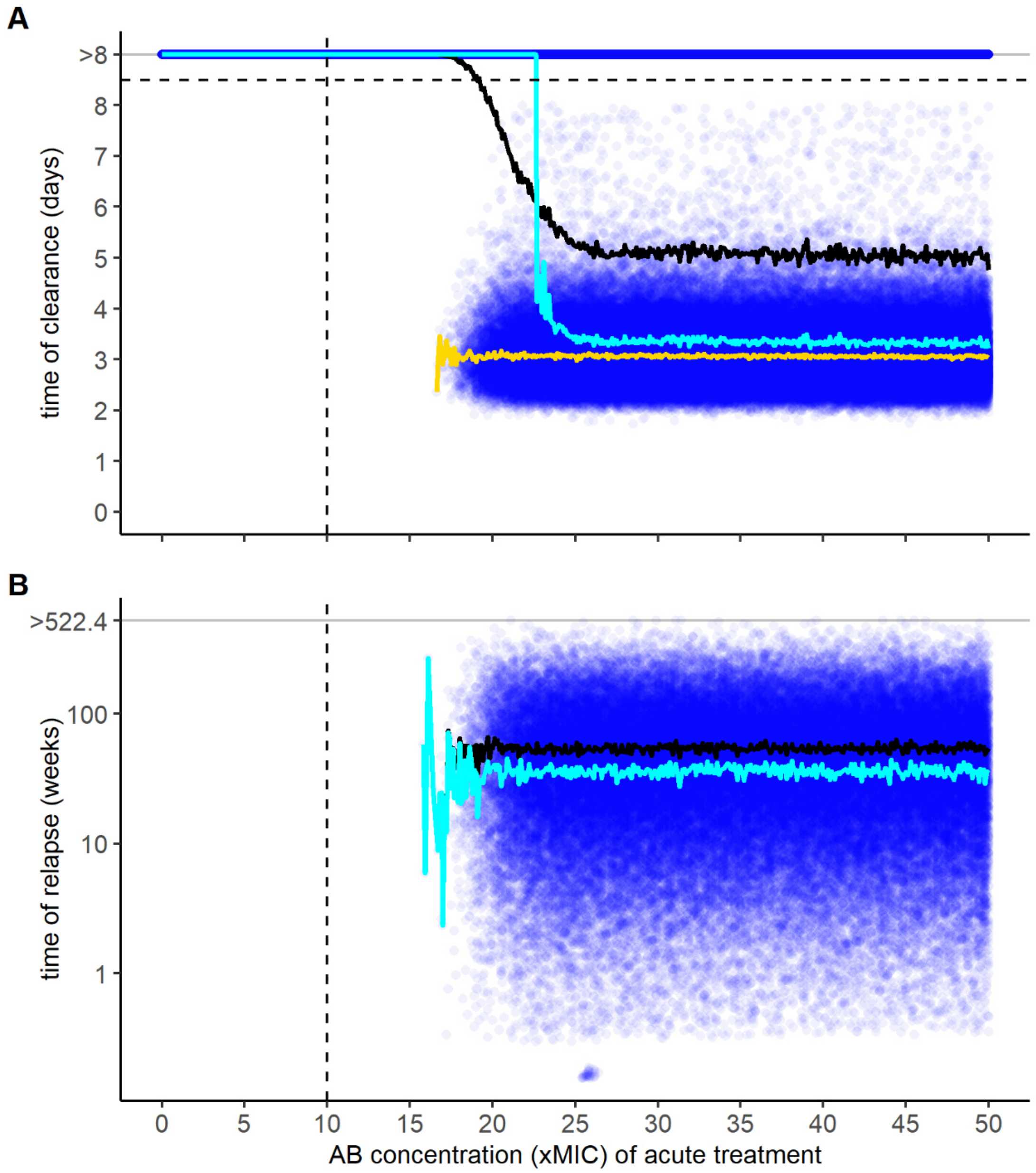


**Figure S7. Time of clearance and relapse. A**) Shown are the time (days) when all bacteria are eradicated for 8 days of acute AB treatment over a range of AB concentrations (0-50xMIC). If no clearance happened during the 8-day treatment period it is depicted as >8 days (line of blue dots). Median (cyan) and mean (black) time of clearance, as well as the mean time of clearance only for simulations where clearance did happen (yellow) are shown. Individual simulation runs are shown as blue dots (1000 runs per AB concentration). **B**) shows the time (weeks) until populations, which survived acute AB treatment (line of blue dots in **A**) at less than $10^5$ CFU start growing and reach $10^6$ CFU over a period of 10 years (522.4 weeks). Median (cyan) and mean (black) time of relapse are shown. Note that the time to relapse is mainly determined by the time it takes surviving persister cells to switch back to the growing state and not by how long it takes for these growing cells to reach $10^6$ CFU.

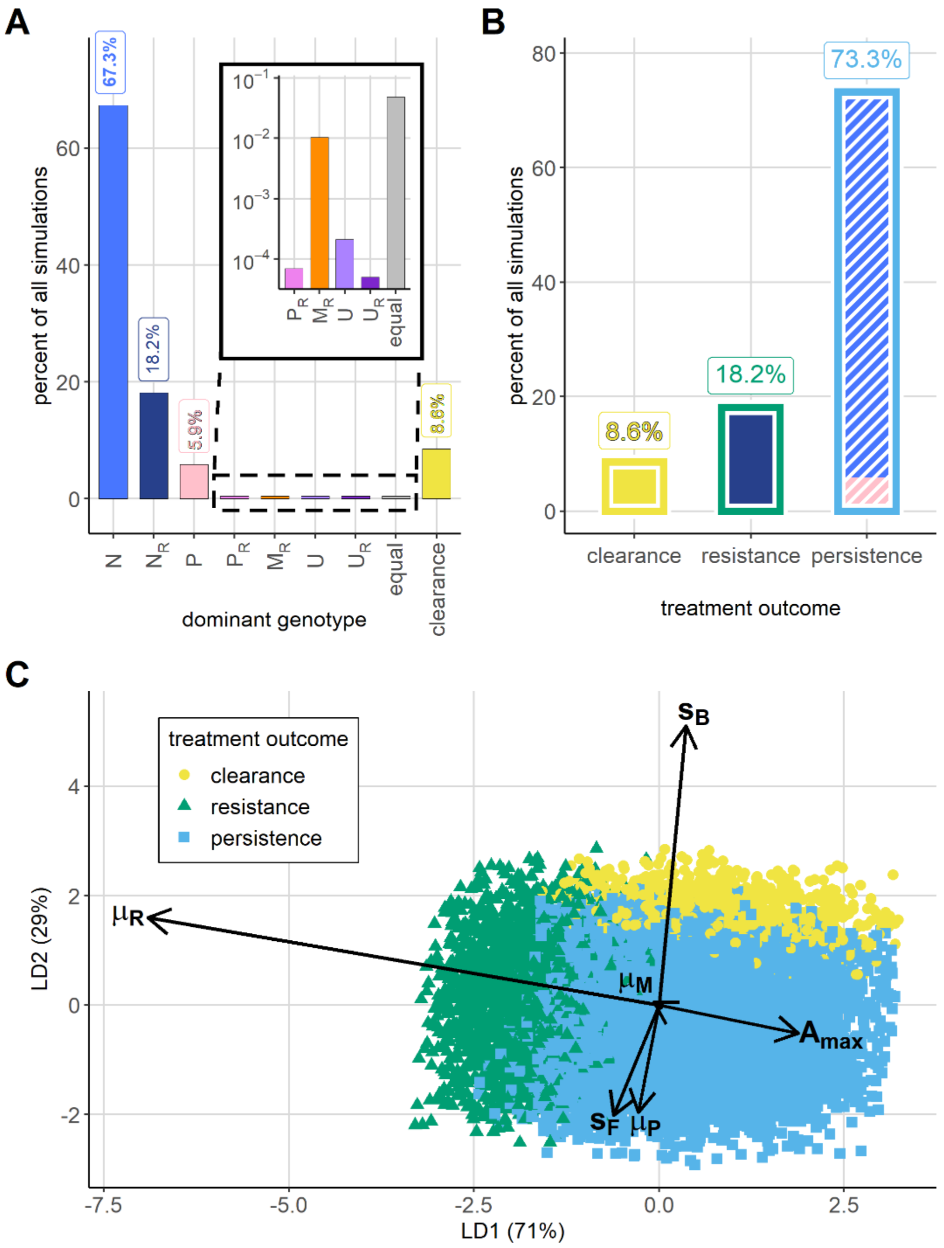


**Figure S8. Parameter sensitivity analysis using random sampling and Linear Discriminant Analysis (LDA). A)** The dominant genotype at the end of 8-day AB treatment shown as percentages of 10,000,000 simulations (100 stochastic simulations each for 10,000 sets of random parameters; see Methods). 'Equal' refers to multiple genotypes being equally abundant (Methods) and 'clearance' to bacterial eradication. The inset shows the genotypes with very small percentages on a log-scale. **B**) Classification of surviving genotypes into treatment outcomes (in % of all simulations) according to Figure 4A. Bar fills show the proportion of dominant genotypes within the classes (colors as in **A**) and bar outlines the class type: yellow shows clearance, green resistance and blue persistence. **C**) LDA for the three treatment outcome classes shown in **B**). Relative lengths and directions of the parameter arrows reflect their direction and magnitude in class separation. For example, $\mu_R$ and $s_B$ display the strongest influence on pushing treatment outcome towards resistance or clearance respectively. Parameter arrows are amplified by a factor of 10 for better visibility. Note that only 0.1% of all data points are shown for clarity.

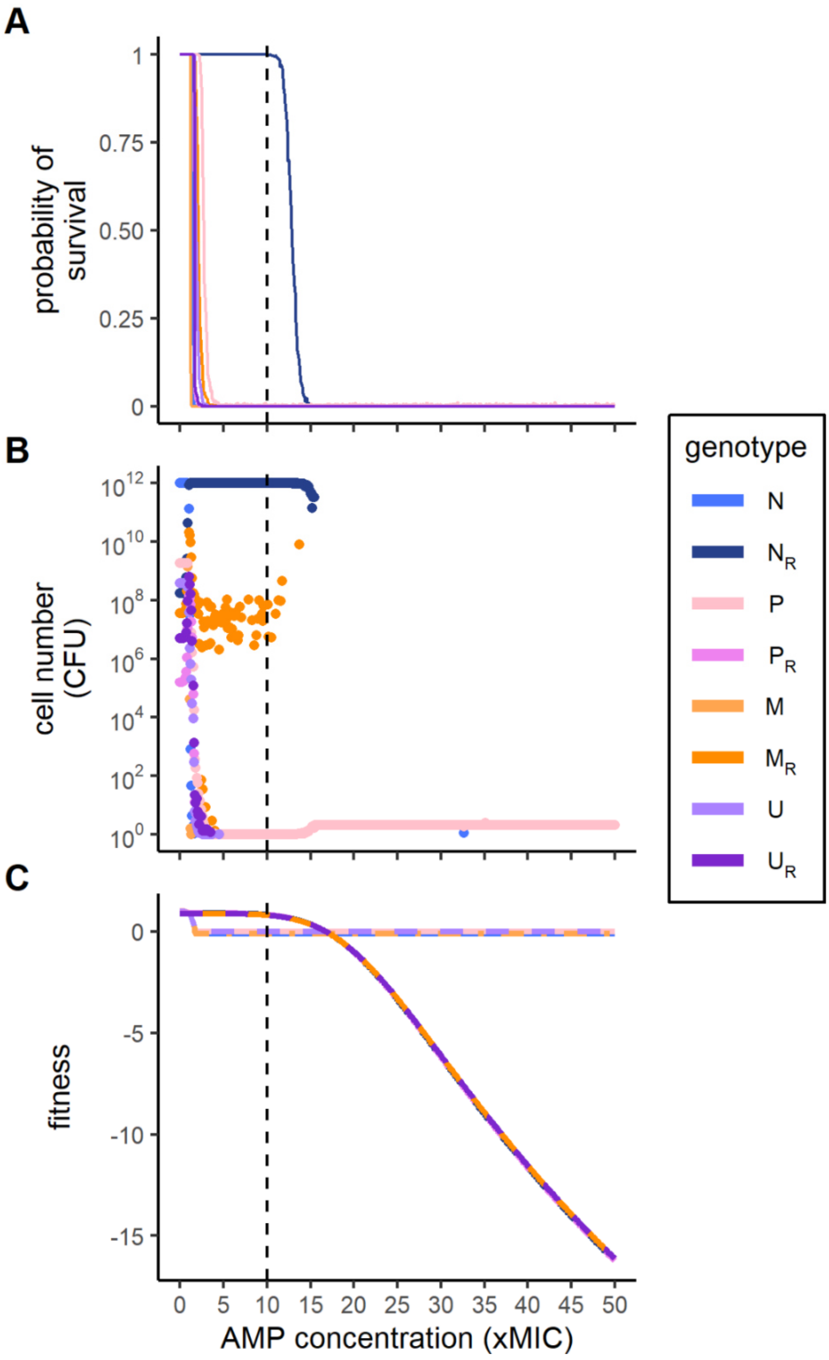


**Figure S9. Effect of Antimicrobial peptide (AMP) concentration on treatment failure due to resistant or persistent genotypes.** Shown are **A**) the probability of genotype survival at the end of treatment, **B**) the mean absolute cell numbers of surviving genotypes, and **C**) the deterministic fitness values of the genotypes. AMPs are small proteins, which display potent antibacterial activity. Note that in **C**) the fitness curve for $N_R$ ($P_R$) largely overlaps with that for $M_R$ ($U_R$) and the fitness curve for N (P) largely overlaps with that for M (U). The pharmacodynamic parameters of AMPs (Table S1) are potentially more favorable for antimicrobial treatments than ABs, as bacteria are killed faster ($\psi_{min}$ = -50) and the dose-response curve is steeper ($\kappa$ = 5), which overall leaves bacteria less opportunity to mutate.

**Table S1. Model parameter values.**

| | Parameter | Description | Value | Unit | Source | Sensitivity analysis range |
|---|---|---|---|---|---|---|
| Pharmacokinetics | $A_{max}$ | peak AB concentration | 0-50 | xMIC | this paper | [15, 30] |
| | $k$ | AB degradation rate | 0.1 | $h^{-1}$ | based on estimates from Igler et al. (2021) and Yu et al. (2018) | - |
| | $\tau$ | time between treatments | 12 | h | this paper | - |
| | $t_{max}$ | maximum run time of acute treatment/relapse simulations | 192/ 87600 | h | this paper | - |
| Pharmacodynamics | $MIC$ | relative Minimal Inhibitory Concentration of susceptibles | 1 | - | this paper | - |
| | $MIC_R$ | relative Minimal Inhibitory Concentration of resistant mutants | 10 | - | this paper | - |
| | $K$ | carrying capacity | $10^{12}$ | CFU | based on estimates from Stressmann et al. (2011) | - |
| | $\psi_{min}$ | minimal net growth with ABs/AMPs | -5/-50 | $h^{-1}$ | Yu et al. (2016) | - |
| | $\psi_{max}$ | maximal net growth without antimicrobials | 1 | $h^{-1}$ | Yu et al. (2016) | - |
| | $\kappa$ | Hill Parameter of AB/AMP dose-response curve | 1.5/5 | - | Yu et al. (2016) | - |
| | $c_R$ | cost of resistance | 0.1 | - | this paper | - |
| | $c_M$ | cost of hypermutation | 0.029 | - | Montanari et al. 2007; Text S5 | - |
| | $d$ | natural death rate of growing cells | 0.01 | $h^{-1}$ | adapted from (Igler et al., 2021) | - |
| Mutation rates | $\mu_R$ | mutation rate from susceptible to resistant | $1x10^{-6}$ | - | Adapted from Yu et al. (2018) | $[10^{-10}, 10^{-5}]$ |
| | $h_\mu$ | fold mutation rate increase for hyper-mutators | 230 | - | Lee et al. 2012 | - |
| | $\mu_M$ | mutation rate of N to M (as multiple of $\mu_R$) | 10x$\mu_R$ | - | this paper; Text S6 | [0.1, 100] |
| | $\mu_P$ | mutation rate of N to P (as multiple of $\mu_R$) | 10x$\mu_R$ | - | this paper; Text S6 | [0.1, 100] |
| Switching | $s_F$ | switching rate into persister state | $1.2x10^{-6}$ | $h^{-1}$ | Balaban et al. 2004 | $[10^{-7}, 1]$ |
| | $s_B$ | back-switching rate from persister to growing state | 0.1 | $h^{-1}$ | Balaban et al. 2004 | $[10^{-7}, 1]$ |
| | $h_F$ | fold increase of switching rate into persister state for high-persisters | 833.3 | - | Balaban et al. 2004 | - |
| | $h_B$ | fold decrease of back-switching rate from persister to growing state for high-persisters | $10^{-3}$ | - | Balaban et al. 2004 | - |

**Table S2. Known mutator genes.**

| Gene | Organism | Source |
|---|---|---|
| *dam* | *E. coli* | Marinus and Morris (1975) |
| *dnaE* | *E. coli* | Maki et al. (1991) |
| *dnaX* | *E. coli* | Henson et al. (1979); Pham et al. (2006) |
| *mutA* | *E. coli* | Michaels et al. (1990) |
| *mutC* | *E. coli* | Michaels et al. (1990) |
| *mutD / dnaQ* | *E. coli* | Degnen and Cox (1974), Horiuchi et al. (1978) |
| *mutH* | *S. enterica* | LeClerc et al. (1996) |
| *mutL* | *E. coli* | Aronshtam and Marinus (1996) |
| *mutM* | *E. coli* | Cabrera et al. (1988) |
| mutS | *E. coli, S. enterica* | LeClerc et al. (1996) |
| *mutT* | *E. coli* | Treffers et al. (1954) |
| *mutY* | *E. coli* | Michaels et al. (1992) |
| *mutY* | *E. coli* | Nghiem et al. (1988) |
| *ndk* | *E. coli* | Lu et al. (1995) |
| *oxyR* | *S. typhimurium* | Storz et al. (1987) |
| *recD* | *E. coli* | Harris et al. (1994) |
| *sodA (sodB)* | *E. coli* | Farr et al. (1986) |
| *ung* | *E. coli* | Duncan and Miller (1980) |
| *uvrD (mutU)* | *S. enterica* | LeClerc et al. (1996) |

**Table S3**. **Persistence-conferring genes in *E. coli.***

| Gene | Source |
|---|---|
| *cspD* | Kim and Wood (2010) |
| *dnaJ* | Vazquez-Laslop et al. (2006) |
| *gadC* | Van den Bergh et al. (2016) |
| *glpD* | Spoering et al. (2006) |
| *hha* | Kim and Wood (2010) |
| *hipA* | Moyed and Bertrand (1983) |
| *hokA* | Kim and Wood (2010) |
| *mazF* | Vazquez-Laslop et al. (2006) |
| *mqsR* | Kim and Wood (2010) |
| *nuoN* | Van den Bergh et al. (2016) |
| *oppB* | Van den Bergh et al. (2016) |
| *phoU* | Li and Zhang (2007) |
| *plsB* | Spoering et al. (2006) |
| *recA* | Dörr et al. (2009) |
| *recB* | Dörr et al. (2009) |
| *relA* | Korch et al. (2003) |
| *relA* | Debbia et al. (2001); Dörr et al. (2009) |
| *relBE* | Keren et al. (2004) |
| *smpB* | Li et al. (2013) |
| *ssrA* | Li et al. (2013) |
| *sucB* | Ma et al. (2010) |
| *tisAB* | Dorr et al. (2010) |
| *tnaA* | Vega et al. (2012) |
| *ubiF* | Ma et al. (2010) |
| *xerC* | Dörr et al. (2009) |
| *xerD* | Dörr et al. (2009) |

**Table S4**. ***E. coli* genes conferring chromosomal AB resistance (via point mutation).** Adapted from Zankari et al. (2017).

| Gene | Antibiotic resistance |
|---|---|
| *pmrA* | colistin |
| *pmrB* | colistin |
| *16S rrsB* | gentamicin |
| *16S rrsC* | kasugamycin |
| *23S* | macrolide |
| *gyrA* | quinolone |
| *gyrB* | quinolone |
| *parC* | quinolone |
| *parE* | quinolone |
| *rpoB* | rifamycin |
| *16S rrsB* | spectinomycin |
| *16S rrsH* | spectinomycin |
| *folP* | sulphonamides |
| *16S rrsB* | tetracycline |

**Text S1. Hyper-mutator frequencies and persister numbers from the literature.**

The frequencies of *Pseudomonas aeruginosa* hyper-mutators from different origins were collected from the literature (Figure 1B): Percentage values reflect the percentage of patients with hyper-mutators. If patient level data was not available, we show the percentage of isolates with hyper-mutators and the study is marked with an asterisk (*). For chronic infections we report the percentage of patients with at least one hyper-mutator strain over the course of infection as it is commonly done. Samples marked as “acute (epidemic)” refer to *Pseudomonas aeruginosa* isolates associated with epidemic outbreaks.

*In vitro* persister numbers (median %) for different antibiotic classes (Figure 1D) are derived from a systematic meta-analysis by Salcedo-Sora and Kell (2020), which collated 187 publications, including 54 antibiotics and 36 bacterial species. We adopted the grouping of individual antibiotics into classes and modes of action used in the original publication. Percentage survival of *in vivo* early and late isolates taken during chronic *Pseudomonas aeruginosa* infections of cystic fibrosis patients are averages over data extracted from Figure 4 of Mulcahy et al. (2010). Mode of action for the antibiotic used by Mulcahy at al. (2010) is given according to the categorization made by Salcedo-Sora and Kell (2020).

**Text S2. Differential equations.** Terms which describe similar mechanisms are colored: maximal growth, effect of AB, persister state switching, growth costs, mutation rates, natural death and carrying capacity.

$$(Eq.\ S1)\ \frac{dN}{dt} = \Big[\psi_{max} * (1 - \mu_R - \mu_M - \mu_P) - \Big(d + (E_N(A(t)) + (1 - \mu_R - \mu_M - \mu_P) * \frac{total}{K} * \psi_{max})\Big) - s_F\Big] * N + s_B * N_p$$

$$(Eq.\ S2)\ \frac{dN_p}{dt} = s_F * N - s_B * N_p$$

$$(Eq.\ S3)\ \frac{dN_R}{dt} = \mu_R * \psi_{max} * N + \Big[\psi_{max} * (1 - c_R) - \Big(d + (E_R(A(t)) + (1 - c_R) * \frac{total}{K} * \psi_{max})\Big)\Big] * N_R$$

$$(Eq.\ S4)\ \frac{dP}{dt} = \mu_P * \psi_{max} * N + \Big[\psi_{max} * (1 - \mu_R - \mu_M) - \Big(d + (E_N(A(t)) + (1 - \mu_R - \mu_M) * \frac{total}{K} * \psi_{max})\Big) - h_F * s_F\Big] * P + h_B * s_B * P_p$$

$$(Eq.\ S5)\ \frac{dP_p}{dt} = h_F * s_F * P - h_B * s_B * P_p$$

$$(Eq.\ S6)\ \frac{dP_R}{dt} = \mu_R * \psi_{max} * P + \Big[\psi_{max} * (1 - c_R) - \Big(d + (E_R(A(t)) + * (1 - c_R) * \frac{total}{K} * \psi_{max})\Big)\Big] * P_R$$

$$(Eq.\ S7)\ \frac{dM}{dt} = \mu_M * \psi_{max} * N + \Big[\psi_{max} * (1 - c_M) * (1 - h_\mu * \mu_R - h_\mu * \mu_P) - \Big(d + (E_M(A(t)) + (1 - h_\mu * \mu_R - h_\mu * \mu_P) * (1 - c_M) * \frac{total}{K} * \psi_{max})\Big) - s_F\Big] * M + s_B * M_p$$

$$(Eq.\ S8)\ \frac{dM_P}{dt} = s_F * M - s_B * M_p$$

$$(Eq.\ S9)\ \frac{dM_R}{dt}$$
$$= h_\mu * \mu_R * \psi_{max} * (1 - c_M) * M$$
$$+ \Big[ \psi_{max} * (1 - c_R) * (1 - c_M)$$
$$- \Big( d + (E_{M_R}(A(t)) + (1 - c_R) * (1 - c_M) * \frac{total}{K} * \psi_{max} \Big) ) \Big] * M_R$$

$$(Eq.\ S10)\ \frac{dU}{dt}$$
$$= h_\mu * \mu_P * \psi_{max} * (1 - c_M) * M + \mu_M * \psi_{max} * P$$
$$+ \Big[ \psi_{max} * (1 - c_M) * \big(1 - h_\mu * \mu_R\big)$$
$$- \Big( d + (E_M(A(t)) + \big(1 - h_\mu * \mu_R\big) * (1 - c_M) * \frac{total}{K} * \psi_{max} ) \Big)$$
$$- h_F * s_F \Big] * U + h_B * s_B * U_p$$

$$(Eq.\ S11)\ \frac{dU_p}{dt} = h_F * s_F * U - h_B * s_B * U_p$$

$$(Eq.\ S12)\ \frac{dU_R}{dt}$$
$$= h_\mu * \mu_R * \psi_{max} * (1 - c_M) * U$$
$$+ \Big[ \psi_{max} + (1 - c_R) * (1 - c_M)$$
$$- \Big( d + (E_{M_R}(A(t)) + (1 - c_R)\ (1 - c_M)\ * \frac{total}{K} * \psi_{max}) \Big) \Big] * U_R$$

**Text S3. Closed net growth integrals of the pharmacodynamic functions over τ time units.**

We include the natural death rate *d* for all genotypes. Note that N and P, $N_R$ and $P_R$, M and U, and $M_R$ and $U_R$, respectively, have the same net growth rate (as determined by the pharmacodynamic function; Eq. 2) and thereby also share the same integrals. Terms which are genotype-characteristic (Table 1) are highlighted in yellow. See Table S1 for descriptions and values of all parameters used.

$$\overline{\Psi}_N = \overline{\Psi}_P = \frac{1}{\tau} * \int_0^{\tau} \Psi_N(t)\, dt$$

$$= \frac{1}{\tau}\left( (\Psi_{max} - d) * \tau + \frac{\log\left(\Psi_{max} * \left(\frac{A_{max}}{\boldsymbol{MIC}}\right)^{\kappa} - \Psi_{min}\right) * (\Psi_{min} - \Psi_{\max})}{k * \kappa} - \frac{log\left(\Psi_{max} * \left(\frac{A_{max} * e^{(-k*\tau)}}{\boldsymbol{MIC}}\right)^{\kappa} - \Psi_{min}\right) * (\Psi_{min} - \Psi_{\max})}{k * \kappa}\right)$$

$$\overline{\Psi}_{N_R} = \overline{\Psi}_{P_R} = \frac{1}{\tau} * \int_0^{\tau} \Psi_{N_R}(t)\, dt$$

$$= \frac{1}{\tau}\left( ((1 - \boldsymbol{c_R}) * \Psi_{max} - d) * \tau + \frac{\log\left((1 - \boldsymbol{c_R}) * \Psi_{max} * \left(\frac{A_{max}}{\boldsymbol{MIC_R}}\right)^{\kappa} - \Psi_{min}\right) * (\Psi_{min} - (1 - \boldsymbol{c_R}) * \Psi_{\max})}{k * \kappa} - \frac{log\left((1 - \boldsymbol{c_R}) * \Psi_{max} * \left(\frac{A_{max} * e^{(-k*\tau)}}{\boldsymbol{MIC_R}}\right)^{\kappa} - \Psi_{min}\right) * (\Psi_{min} - (1 - \boldsymbol{c_R}) * \Psi_{\max})}{k * \kappa}\right)$$

$$\overline{\Psi}_M = \overline{\Psi}_U = \frac{1}{\tau} * \int_0^{\tau} \Psi_M(t)\, dt =$$

$$= \frac{1}{\tau}\Bigg( ((1 - \boldsymbol{c_M}) * \Psi_{max} - d) * \tau$$

$$+ \frac{\log\left((1 - \boldsymbol{c_M}) * \Psi_{max} * \left(\frac{A_{max}}{\boldsymbol{MIC}}\right)^{\kappa} - \Psi_{min}\right) * (\Psi_{min} - (1 - \boldsymbol{c_M}) * \Psi_{\max})}{k * \kappa}$$

$$- \frac{log\left((1 - \boldsymbol{c_M}) * \Psi_{max} * \left(\frac{A_{max} * e^{(-k*\tau)}}{\boldsymbol{MIC}}\right)^{\kappa} - \Psi_{min}\right) * (\Psi_{min} - (1 - \boldsymbol{c_M}) * \Psi_{\max})}{k * \kappa}\Bigg)$$

$$\overline{\Psi}_{M_R} = \overline{\Psi}_{U_R} = \frac{1}{\tau} * \int_0^{\tau} \Psi_{M_R}(t)\, dt =$$

$$= \frac{1}{\tau}\Bigg( ((1 - \boldsymbol{c_M}) * (1 - \boldsymbol{c_R}) * \Psi_{max} - d) * \tau$$

$$+ \frac{\log\left((1 - \boldsymbol{c_M}) * (1 - \boldsymbol{c_R}) * \Psi_{max} * \left(\frac{A_{max}}{\boldsymbol{MIC_R}}\right)^{\kappa} - \Psi_{min}\right) * (\Psi_{min} - (1 - \boldsymbol{c_M}) * (1 - \boldsymbol{c_R}) * \Psi_{\max})}{k * \kappa}$$

$$- \frac{log\left((1 - \boldsymbol{c_M}) * (1 - \boldsymbol{c_R}) * \Psi_{max} * \left(\frac{A_{max} * e^{(-k*\tau)}}{\boldsymbol{MIC_R}}\right)^{\kappa} - \Psi_{min}\right) * (\Psi_{min} - (1 - \boldsymbol{c_M}) * (1 - \boldsymbol{c_R}) * \Psi_{\max})}{k * \kappa}\Bigg)$$

**Text S4. The impact of timing on U emergence and establishment.**

The timing of U emergence could be crucial for its survival given that the AB concentration changes over time due to degradation and periodic administration of AB doses. The stochastic switching into the persister state – which makes high-persistence beneficial – is less likely in the presence of high AB concentrations, because bacteria are killed quickly, giving them little time to switch into persister state. Further, switching back to a growing state will kill non-resistant M, P, and U populations. Therefore, the chance for U to emerge from P and survive by switching into the persister state is higher in the timeframe where AB concentrations are low. Consequently, the larger persister population of P could facilitate the emergence and establishment of U via the persister state more often than M.

**Text S5. Determination of hyper-mutator growth costs.**

Montanari et al. (2007) report doubling times for seven hypermutable strains of *Pseudomonas aeruginosa* isolated from cystic fibrosis patients. We transform these doubling times (DT) to growth rates (GR) by using the formula $GR = \frac{1}{DT} * \log(2)$. Generally, we assume that a hyper-mutator mutation is – per se – accompanied by a growth cost and that the accumulation of negative and neutral mutations outweighs the effects of positive mutations. While this may overestimate the immediate negative effects of hyper-mutation and underestimate the effects of beneficial mutations over time, we are interested here only in investigating the impact of hyper-mutation on facilitation of specific mutations. Of the seven strains from Montanari et al. (2007), we therefore disregard all strains, which after complementation of their hyper-mutation-causing genetic changes (i.e. functional genes re-introduced via plasmids) display equal or higher growth rates than the WT as this could be due to the accumulation of beneficial mutations. This leaves a *mutS* mutant (BT1) and a *mutL* mutant (RP74), for which we determine the inherent growth cost of hypermutation by calculating the relative difference in growth rates of hyper-mutator strains and the complemented mutators. The respective relative growth costs are 0.049 and 0.009 for *mutS* and *mutL* respectively. For our simulations we use the mean of these two values, 0.029.

**Text S6. Estimation of mutation rates to high-persister and hyper-mutator genotypes.**

While multiple genes involved in hyper-mutation (see Miller, 1996; Horst et al., 1999; Jayaraman, 2009) and persistence (see Zhang, 2014; Kaldalu et al., 2020) are known, the probability that a chromosomal mutation results in a high-persister or hyper-mutator phenotype is unknown. In Table S2 and Table S3 we compiled known mutator genes and genes involved in persistence of *E. coli* and other pathogens from the literature. In total we found 19 mutator genes and 26 persistence genes and hence potential targets for hyper-mutator and high-persistence mutations. In contrast, Zankari et al. (2017) reported that relatively few genes can confer antibiotic resistance via chromosomal point mutations (Table S4). On average 1.4 genes confer resistance against a single AB. Therefore, we estimate mutation rates to high-persisters and hyper-mutators to be at least 10 times higher than the resistance mutation rate, which, considering the number of potential targets, should be a relatively conservative estimate. We vary this parameter however in the LDA to test the sensitivity of our results to this assumption.
